# IL-23 tunes inflammatory functions of human mucosal-associated invariant T (MAIT) cells

**DOI:** 10.1101/2024.08.29.609178

**Authors:** Laetitia Camard, Tharshana Stephen, Hanane Yahia-Cherbal, Vincent Guillemot, Sébastien Mella, Victoire Baillet, Hélène Lopez-Maestre, Daniele Capocefalo, Laura Cantini, Claire Leloup, Julie Marsande, Juan Sienes-Bailo, Ambre Dangien, Natalia Pietrosemoli, Milena Hasan, Huimeng Wang, Sidonia B. G. Eckle, Anne M. Fourie, Carrie Greving, Barbara Joyce Shaikh, Raphaelle Parker, Daniel J. Cua, Elisabetta Bianchi, Lars Rogge

## Abstract

IL-23 signaling plays a key role in the pathogenesis of chronic inflammatory and infectious diseases, yet the cellular targets and signaling pathways affected by this cytokine remain poorly understood. We show that IL-23 receptors are expressed on the large majority of human MAIT, but not of conventional T cells. Protein and transcriptional profiling at the population and single cell level demonstrates that stimulation with IL-23 or the structurally related cytokine IL-12 drives distinct functional profiles, revealing a high level of plasticity of MAIT cells. IL-23, in particular, affects key molecules and pathways related to autoimmunity and cytotoxic functions. Integrated analysis of transcriptomic and chromatin accessibility, supported by CRISPR/Cas9 mediated deletion, shows that AP-1 transcription factors constitute a key regulatory node of the IL-23 pathway in MAIT cells. In conclusion, our findings indicate that MAIT cells are key mediators of IL-23 functions in immunity to infections and chronic inflammatory diseases.

## Introduction

IL-23 plays critical roles in the pathogenesis of both infectious and chronic inflammatory diseases (CID), as demonstrated by the susceptibility of IL-23R-deficient patients to mycobacterial infections, and the success of IL-23-blockers in the treatment of several CID, respectively ^1–3^. Key evidence of the importance of IL-23 in brain, joint and intestinal inflammation came from the analysis of mice deficient for IL-23 or the IL-23 receptor (IL-23R) ^4–8^.

In parallel, genome-wide association studies (GWAS) revealed a strong association of genetic variants in the *IL23R* gene and other molecules in this signaling pathway with Crohn’s disease (CD), psoriasis (Pso) and axial spondyloarthritis (axSpA) ^9,10^. *IL23R* variants affect both IL-17A and IFNγ production by human CD4^+^ T cells from axSpA patients, and stimulation of activated CD4^+^ T cells with IL-23 increased IL-17A and IFNγ secretion ^11,12^.

Experimental models have shown that IL-23 plays a central role for the expansion and function of proinflammatory T helper 17 (Th17) cells ^8,13^. In addition, more recent studies have pointed to important roles of innate-like immune cells expressing the IL-23 receptor (IL-23R) in the pathogenesis of CID. Systemic expression of IL-23 induced hallmarks of SpA in a mouse model mediated by CD3^+^CD4^-^CD8^-^RORγt^+^ cells, establishing a direct link between IL-23 and tissue inflammation ^14^. IL-23R^+^ γδ T cells were enriched in the blood of SpA patients ^15^ and IL-23 was shown to increase production of IL-17 and IL-21 by RORγt^+^ iNKT and γδ T cells ^16^. Mucosal-associated invariant T (MAIT) cells from SpA patients were also shown to produce IL-17 ^17^ and we have recently demonstrated that MAIT cells from axSpA patients expressed the highest levels of *IL23R* transcripts ^18^.

MAIT cells are innate-like T cells that express a semi-invariant Vα7.2-Jα33/12/20 (*TRAV1-2 – TRAJ33/12/20*) T cell receptor (TCR) ^19,20^. These cells predominantly reside in tissues, such as the liver, lung, skin and the gastrointestinal tract ^21^. MAIT cells are restricted by the MHC class I-related protein 1 (MR1) presenting riboflavin (vitamin B2) metabolites ^22,23^. These cells have important functions in anti-microbial immunity but also in tissue repair and homeostasis ^24–28^. In addition to MR1-bound antigens, MAIT cells can be activated by cytokines, in particular IL-12 and IL-18, but also by type I IFN, IL-7, IL-15 and TNF, underlining their innate-like characteristics ^17,27,29^. A recent study has shown that patients homozygous for loss-of-function *IL23R* alleles display Mendelian susceptibility to mycobacterial diseases (MSMD) and, less frequently, chronic mucocutaneous candidiasis (CMC). In this context, IL-23 is necessary for *Mycobacterium*-induced type 1 immune responses and *Candida*-induced type 17 immunity in MAIT cells, documenting a non-redundant role of IL-23 signaling in this cell population ^2^.

However, little is known about the signaling pathways and regulatory networks affected by IL-23 in human MAIT cells. Here, we have compared signaling by IL-23 and the structurally related cytokine IL-12 by protein and transcriptional profiling at the population and single-cell level of circulating human MAIT cells. Integration of these data with the analysis of open chromatin revealed a remarkable level of plasticity of these cells and pointed to the AP-1 transcription factor BATF as a key regulatory node affecting MAIT cell functions.

## Results

### A large majority of human MAIT cells express IL-23R on their surface

IL-23 plays key roles in infectious and inflammatory diseases ^2,10^, therefore the identification of the immune cells that respond to IL-23 is important for the understanding of the pathophysiology of these diseases. However, monitoring the expression of the receptor for IL-23 on human cells has remained challenging because of the inadequate specificity and/or sensitivity of commercially available reagents. Using a newly generated anti-IL-23R monoclonal antibody, we found that MAIT cells from peripheral blood of healthy donors were the population with the highest frequency of IL-23R expressing cells (median: 62.21%, interquartile range (IQR): 48.65 – 71.66), followed by CD3^+^CD56^+^ cells (18.03%, 8.96 – 13.1) and γο T cells (12.16%, 7.25 – 18.76). In contrast, only low percentages of resting CD4^+^ and CD8^+^ T cells or NK cells expressed IL-23R (1.89, 1.82 and 1.54% respectively) (**Figure 1A**, **Table S1**). These data are consistent with our previous analysis of *IL23R* transcripts in stimulated MAIT, γδ, CD4^+^ and CD8^+^ T cells isolated from PBMC of axial spondyloarthritis (axSpA) patients ^18^.

**Fig. 1.**
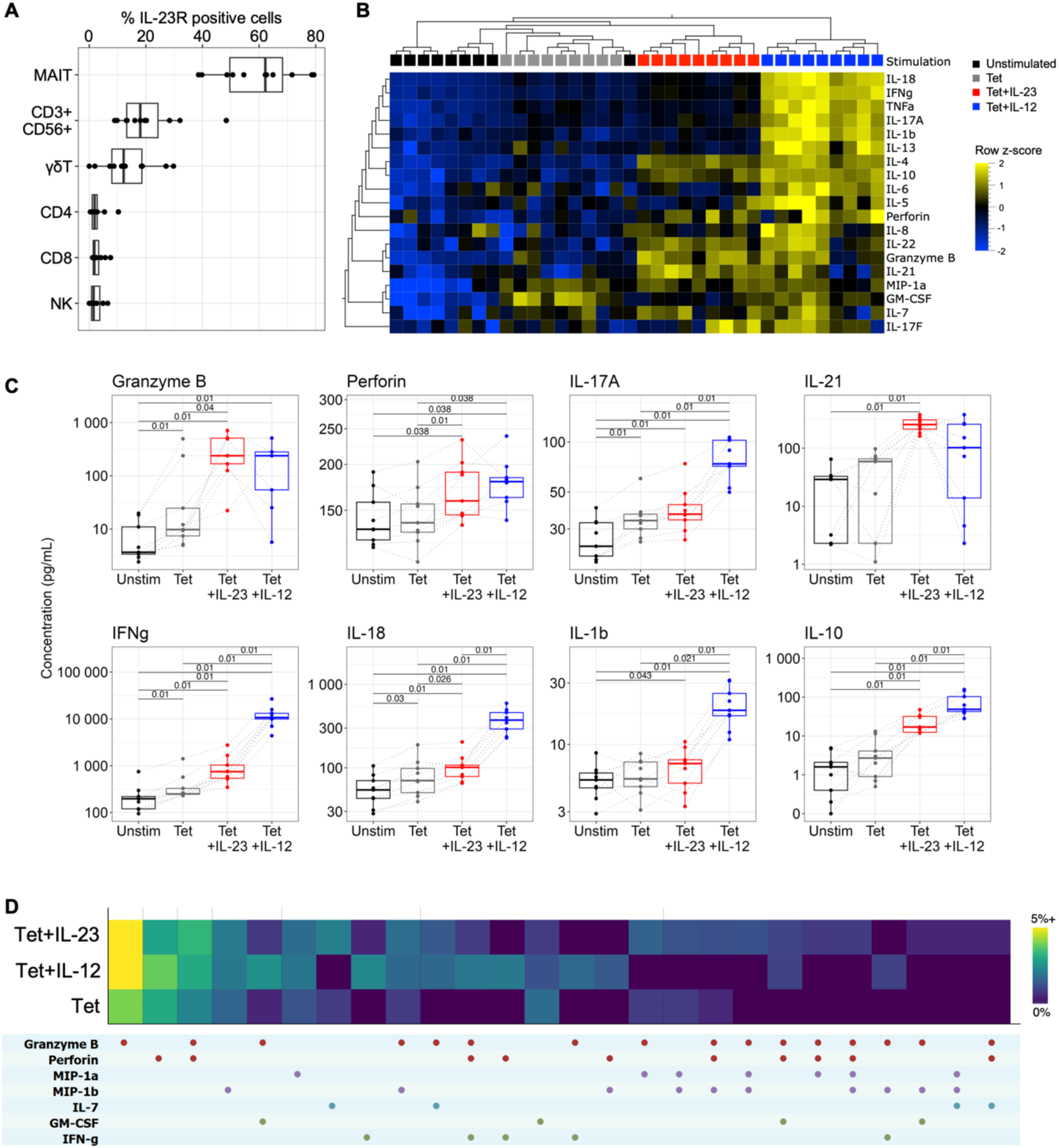
IL-23 enhances production of cytotoxic molecules and IL-10 by MAIT cells. **(A)** Percentage of IL-23R positive cells in different immune cell populations was assessed by flow cytometry on freshly isolated PBMCs (n=11). (**B** and **C**) PBMC were activated with Tet or left unstimulated (black) for 6 days. Sorted MAIT cells (CD3^+^CD161^+^Va7.2^+^) were re-stimulated with Tet in the absence (grey) or presence of IL-23 (red) or IL-12 (blue) for 24h. Cytokine secretion in the supernatants was measured using Luminex technology (n=9). (**B**) Heatmap of measured cytokines. Dendograms on the top and left sides correspond to hierarchical clustering. (**C**) Boxplots representing the concentration of the indicated cytokines (adjusted p-values, paired Wilcoxon test, Benjamini-Hochberg correction). (**D**) Single cell secretome of MAIT cells activated for 6 days with Tet in the presence or absence of IL-23 or IL-12 was assessed using Isolight technology. Heatmap represents the single-cell co-secretion patterns and their frequencies (n=4).

Given the preferential expression of IL-23R on MAIT cells, we next investigated the function of IL-23 signaling in this population. We stimulated MAIT cells with MR1 tetramers loaded with 5-(2-oxopropylideneamino)-6-D-ribitylaminouracil (MR1/5-OP-RU) ^30^ in the absence or presence of IL-23, or of the structurally related cytokine IL-12. We found that IL-23 significantly enhanced secretion of granzyme B, perforin, IL-21, IFN-γ, IL-18, and IL-10 by MAIT cells (**Figures 1B and C**, **Table S2**), Of note, IL-23 only modestly increased IL-17A production, while secretion of IL-17A, as well as of IL-1β and IL-18 was strongly enhanced in the presence of IL-12 (**Figures 1B and C**). Analysis of protein secretion at the single-cell level showed that IL-23 increased the frequency of MAIT cells producing cytotoxic molecules, compared to the MR1/5-OP-RU tetramer (Tet) alone stimulation (**Figure 1D**). Comparable results were obtained in the presence of IL-12. At the single-cell level we could also detect a small increase in the frequency of MAIT cells secreting IL-7, MIP-1α and MIP-1β in response to IL-23. Most of the cells showed co-expression of different cytokines and cytotoxic molecules, arguing for a polyfunctional response to IL-23 (**Figure 1D**). Together, our analysis of protein secretion both at the bulk and single-cell levels revealed substantially enhanced cytotoxic functions of MAIT cells by IL-23.

### Single-cell profiling reveals the plasticity of MAIT cells

To investigate the heterogeneity of the response to IL-23 and IL-12, we performed “Cellular Indexing of transcriptomes and Epitopes by sequencing” (CITE-seq) ^31^ on MAIT cells from 3 healthy donors, stimulated either with Tet alone (17594 cells after filtering), or Tet+IL-23 (19153 cells) or Tet+IL-12 (9876 cells). We identified 16 clusters (**Figure 2A**) characterized by preferential expression of both genes (**Figure 2B**, **Table S3**) and cell surface markers (**Figures S1A, Table S4).** Projection of the stimulation conditions on the UMAP revealed that cells stimulated with Tet alone or in the presence of IL-23 cluster together (**Figures 2C-D** and **Figure S1B**); while Tet+IL-12 stimulated cells cluster separately, in clusters 2, 11, 12 and 15. Of note, clusters 4 and 5 are composed of cells from the 3 stimulation conditions and correspond to cycling cells (**Figures 2D and S1D**).

**Fig. 2.**
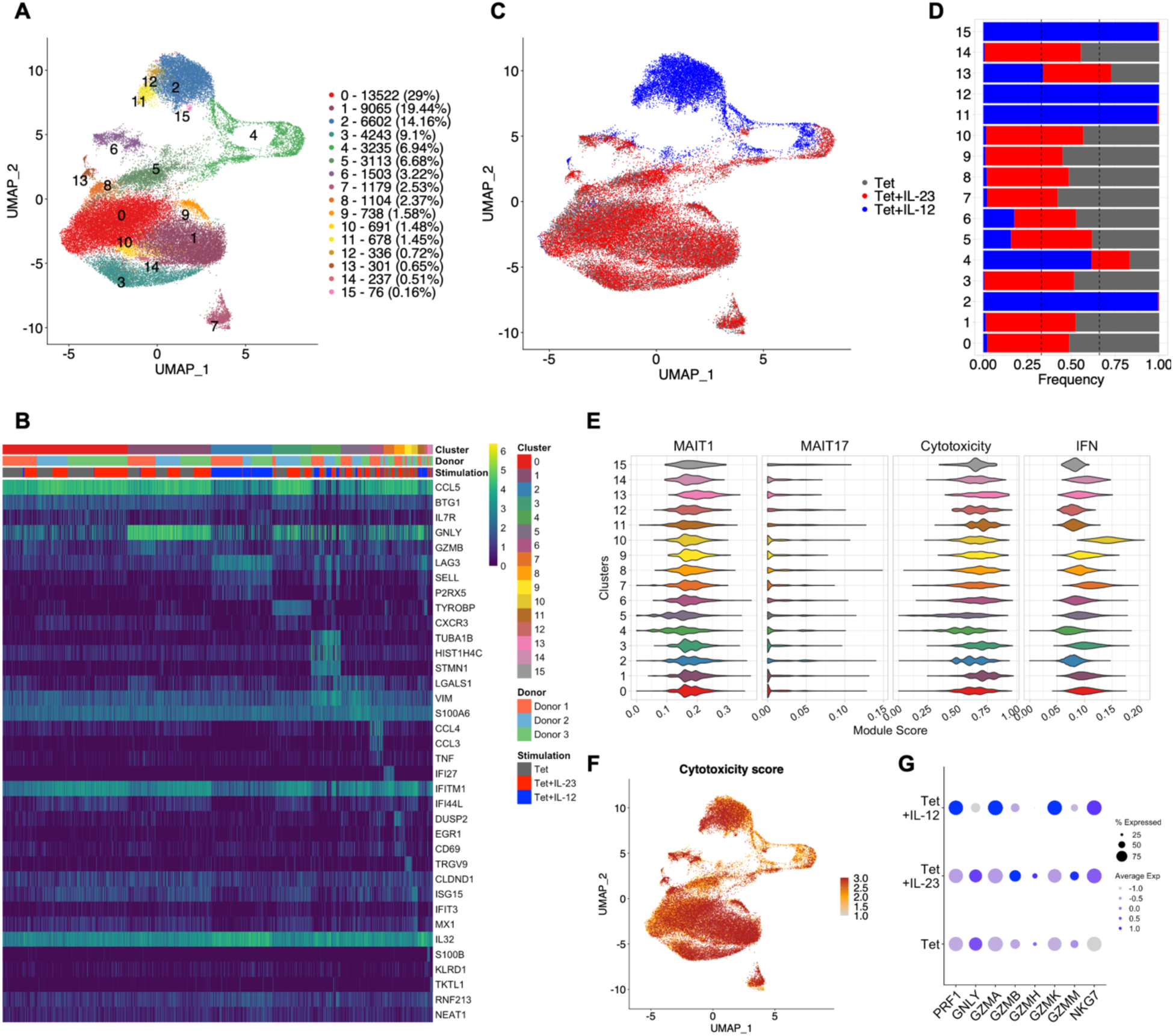
Combined transcriptomic and phenotypic analysis of MAIT cells by single-cell profiling. PBMCs were activated with Tet in the presence or absence of IL-23 or IL-12 for 6 days, MAIT cells were sorted (CD3^+^MR1/5-OP-RU^+^) and CITE-seq was performed (n=3). (**A**) UMAP of stimulated MAIT cells colored by the 16 identified clusters. Numbers and frequencies of cells in each cluster are indicated. (**B**) Heatmap showing row-scaled log-transformed normalized expression of the top marker genes for each MAIT cell cluster. (**C**) UMAP of stimulated MAIT cells colored by the stimulation condition. (**D**) Proportion of cells from the different stimulation groups in each cluster. Grey: Tet, red: Tet+IL-23, blue: Tet+IL-12. (**E**) Violin plots of the scores of the indicated signatures by cluster. (**F**) UMAP of stimulated MAIT cells colored by cytotoxicity genes score. (**G**) Expression of cytotoxicity genes in MAIT cells from the different stimulation groups.

Cluster 2 is characterized by *SELL* (encoding L-selectin, CD62L) and LAG3 (encoding CD223) expression, both at the level of gene expression (**Figures 2B** and **S1C**) and cell surface protein expression (**Figure S1A**). These markers are not restricted to cluster 2, but more globally to IL-12 stimulated cells (**Figure S1C**). CCL3 and CCL4 (encoding MIP-1α and MIP-1β, respectively) characterize cluster 6 (**Figure 2B** and **Figure S1C**).

We identified clusters characterized by the expression of *GNLY*, encoding the cytotoxic molecule granulysin, both in Tet/Tet+IL-23 (cluster 1) and in Tet+IL-12 cells (cluster 11, **Figure 2B** and **Figure S1C**). Similarly, clusters 3 and 12 are characterized by expression of *TYROBP* (encoding DAP12, **Figure 2B** and **Figure S1C**), a signaling molecule containing an ITAM motif that associates with activating receptors, such as KIR2DS2 or CD94/KLRD1 on NK cells ^32,33^.

Previous studies have suggested the presence of distinct MAIT1 and MAIT17 subsets, in particular in the mouse ^34,35^. We asked whether we could identify these subsets in our samples computing UCell scores ^36^ using previously published MAIT1 and MAIT17 signatures, as well as Th1 and Th17 signatures ^37,38^. Consistent with recent reports in humans ^29,37^, our analysis did not reveal specific MAIT1 and MAIT17 subsets (**Figure 2E** and **fig S2A-C**). The apparent increase in the MAIT17 signature in the Tet+IL-12 condition was driven by the increased expression of two genes, namely *ITGAE* (a resident T cell marker) and *IL23R* (**Figure S2B**). Consistent with this notion, recent work has demonstrated that *IL23R* is an IL-12 target gene in murine CD4^+^ T cells ^39^.

Although we did not detect a discrete MAIT1 subset in non-polarizing Tet only conditions, treatment with IL-12 had a marked effect on the expression of genes detectable at the single-cell level, resulting in a functional polarization towards a type 1 response, with increased expression of *TBX21*, *IL12RB2* and *IFNG*, and reduction of *RORC* and *CCR6* expression (**Figure S2B** and **Figure S3**).

With the exception of cycling cells (clusters 4 and 5), all MAIT cells were characterized by a high cytotoxicity genes score (**Figure 2E-F** and **Figure S2C**), however IL-12 and IL-23 stimulated cells did not express the same cytotoxic molecules. IL-12 stimulated MAIT cells expressed preferentially *PRF1*, *GZMA* and *GZMK*, while IL-23 stimulated cells expressed *GNLY*, *GZMB* and *GZMM* (**Figure 2G** and **Figure S4**).

IFN-inducible genes are markers for clusters 7 (*IFI27, IFITM1 and IFI44L*) and 10 (*ISG15 and IFIT3*), suggesting an important role for type I IFN on MAIT cell functions in these specific clusters (**Figure 2B** and **Figure S1C**). Consistently, a high IFN response score was restricted to these two clusters (IFN signature from https://github.com/carmonalab/SignatuR) **Figure 2E**). Cluster 7 specifically expressed high levels of IFI27, while cells in cluster 10 expressed high levels of several IFN inducible genes (**Figure S2D**). The IFN response score was reduced in MAIT cells stimulated in the presence of IL-12 (**Figure S2C**).

Overall, our single-cell profiling of human MAIT cells isolated from the peripheral compartment did not reveal discrete MAIT1 and MAIT17 sub-populations as observed in the mouse, but rather a functional polarization that may reflect the plasticity of these cells.

### Integrative analysis of CITE-seq data reveals IL-23-dependent upregulation of MHC class II genes in MAIT cells

As the analysis described above did not highlight clusters of cells specifically associated to IL-23 stimulation, we performed an additional matrix factorization decomposition of the CITE-seq data. The aim of this matrix factorization step is to jointly reduce the surface protein and gene expression data from CITE-seq into a linear combination of latent factors shared among the two omics datasets. Each latent factor represents a biological signal contributing to the state of the cells under analysis ^40^. We used Multi-Omics Wasserstein inteGrative anaLysIs (Mowgli), a matrix factorization tool combining Optimal Transport and Non-Negative Matrix Factorization, shown to have higher interpretability than other state-of-the-art tools ^41^. We applied the Mowgli framework to the CITE-seq data keeping the three available donors separated to avoid the batch correction step hiding part of the relevant biological signals.

The UMAPs of the cell embeddings resulting from the integration are provided in **Figure 3A-C**. Based on the UMAP representations, all donors seem to present a separated cluster of cycling cells, in agreement with the previous clustering analysis. In addition, all three donors show some separation between the Tet+IL-12 stimulated cells and the other conditions. Interestingly, the cell embeddings of Donor 1 (**Figure 3A**) also shows some separation between the Tet+IL-23 and Tet conditions. Regarding then the latent factors identified by Mowgli, factors specifically associated to IL-12 and IL-23 stimulations could be identified. Among the factors associated with Tet+IL-23, one was consistently present in all three donors (**Figure 3D-F**) with correlation between its projections on the gene space of ∼0.8 and on the protein space of ∼0.9 (**Figure 3G**). The top contributing genes and proteins associated to the IL-23-specific factor include a striking number of the MHC class II genes *HLA-DRA*, *HLA-DRB1*, *HLA-DQB1*, *HLA-DMA*, *HLA-DPB1*, *HLA-DPA1*, the genes encoding cytotoxic molecules, such as *GZMH* and *GZMB* (**Figure 3H**), and the anti-HLA-DR surface protein (**Figure 3I**).

**Fig. 3.**
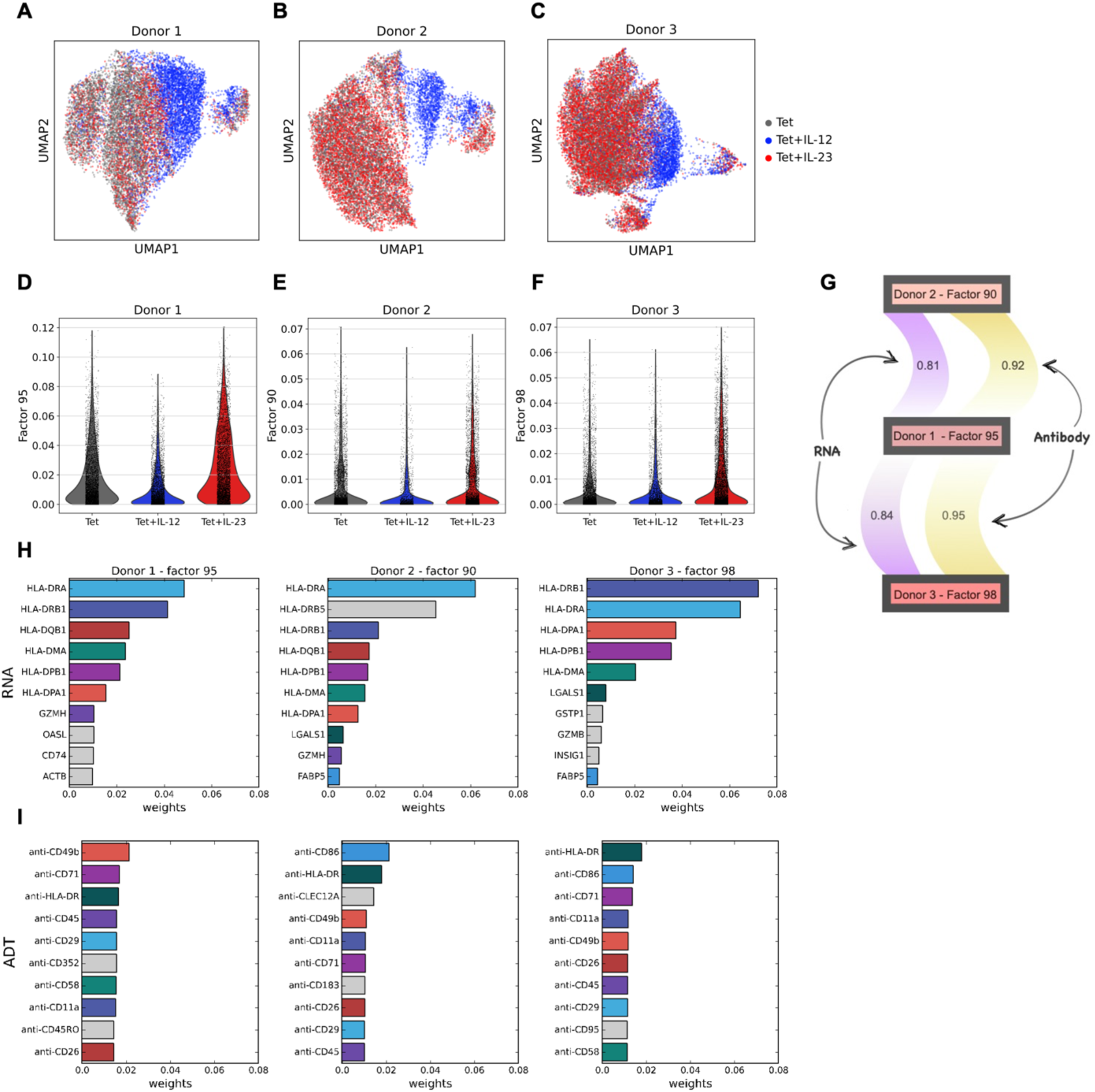
Mowgli analysis of CITE-Seq data identifies factors associated to IL-23 stimulation. **(A to C)** UMAPs of the integration of protein and RNA modalities in CITE-Seq Data for donors 1 (A), 2 (B) and 3 (C). Grey: cells treated with Tet; red: Tet+IL-23; blue: Tet+IL-12 (**D to F)** Distribution of the weights associated to each factor according to treatment. D: Donor 1, Factor 95; E: Donor 2, Factor 90; F: Donor 3, Factor 98. (**G)** Spearman correlation across the factors associated to Tet+IL-23 for each donor (black rectangles) in the RNA (left) or antibody space (right). Correlation are shown only between Donor 1 and Donor 2 or 3. For RNA, only genes in common across the highly variable features of each sample were used to compute the correlation. **(H-I)** Top ten genes (H) and antibodies (I) associated to Tet+IL-23 response in the relevant factor identified by Mowgli in each donor. Colored bars denote the presence of a gene or antibody in the other top ten genes of another donor. Grey bars represent genes or antibody specific for that donor.

Together, this analysis revealed that stimulation of MAIT cells in the presence of IL-23 results in a remarkable regulation of MHC class II and cytotoxic molecules.

### Genes and pathways regulated by IL-23 or IL-12 in MAIT cells

Although the majority of MAIT cells express the IL-23R, very little is known about the genes and signaling pathways regulated by IL-23 in these cells.

We defined transcriptomic signatures of stimulated MAIT cells by RNA sequencing. Stimulation in the presence of IL-23 resulted in differential expression of 607 genes, compared to Tet alone (adj.p<0.05, **Table S5**). Consistent with the single-cell data, stimulation with Tet+IL-12 impacted a larger number of genes (1013, adj.p<0.05, **Table S6**). Analysis of IL-23-versus IL-12-regulated gene expression (**Figure 4A**), revealed that a fraction of these genes (242) were co-regulated by the two cytokines, while 365 and 771 were specifically regulated by IL-23 and IL-12, respectively (**Figure 4B**). Pathway analysis of differentially expressed genes underscored the distinct biological functions of the two cytokines. Genes regulated by IL-23 were enriched in signaling pathways associated with “classical” autoimmune diseases, such as multiple sclerosis (MS), systemic lupus erythematosus (SLE), type 1 diabetes (T1D) and rheumatoid arthritis (RA) (**Figure 4C**, **Table S7**). IL-12-regulated genes, in contrast, were associated with infectious disease pathways, consistent with the role of type 1 responses in these diseases (**Figure 4D**, **Table S8**).

**Fig. 4.**
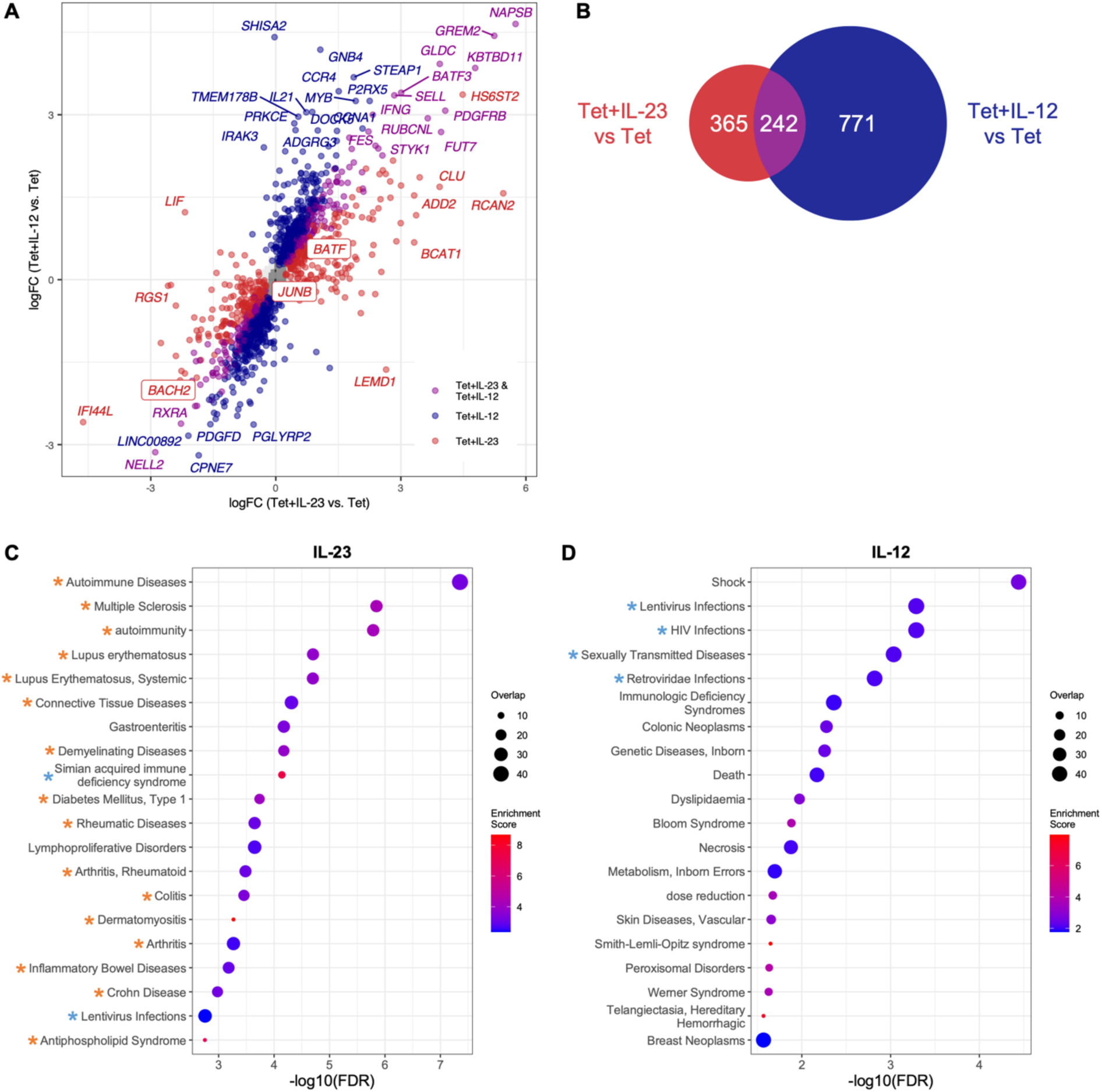
Genes and pathways regulated by IL-23 or IL-12 in MAIT cells. PBMCs were activated with Tet in the presence or absence of IL-23 or IL-12 for 6 days, CD3^+^CD161^+^Va7.2^+^ MAIT cells were sorted, and RNA-sequencing was performed. (n=5-8) (**A**) Scatter plot of changes in gene expression (log2-fold change, x-axis: Tet+IL-23 vs. Tet, y-axis: Tet+IL-12 vs. Tet). Each dot corresponds to a differentially expressed gene (red: Tet+IL-23 vs. Tet, blue: Tet+IL-12 vs. Tet, purple: Tet+IL-23 vs. Tet and Tet+IL-12 vs. Tet). Grey shading indicates the 2D density plot of all the delta log-fold change. (**B**) Euler diagram representing the differentially express genes in the Tet+IL-23 vs. Tet, and in the Tet+IL-12 vs. Tet stimulation conditions. (**C** and **D**) Over-representation analysis was performed on the genes regulated by (**C**) IL-23 or (**D**) IL-12 using the GLAD4U database. Bubble plots of the top 20 associated pathways. Orange asterisks indicate pathways associated with chronic inflammatory and autoimmune diseases pathways. Blue asterisks indicate pathways associated with infectious diseases.

### IL-23 regulates MHC class II and AP-1 transcription factor genes in MAIT cells

Chronic inflammatory diseases associated with genetic variants in the IL-23 signaling pathway, such as CD, Pso and axial SpA ^10^ are primarily linked to innate immune activation and MHC class I genes ^42^. Our observation that IL-23-regulated genes were associated with pathways linked to classical autoimmune diseases was therefore somewhat unexpected as these diseases are typically associated with the adaptive arm of the immune system. We therefore investigated further the expression of the genes enriched in the autoimmune pathways (**Table S7**) in MAIT cells stimulated in the presence of IL-23, and asked how they were affected by IL-12 or IL-23 stimulation (**Figure 5A**). We found that expression of 10 genes encoding MHC class II molecules was upregulated by IL-23 but not by IL-12 (**Figure 5A and B**), contributing to the strong enrichment of autoimmune pathways.

**Fig. 5.**
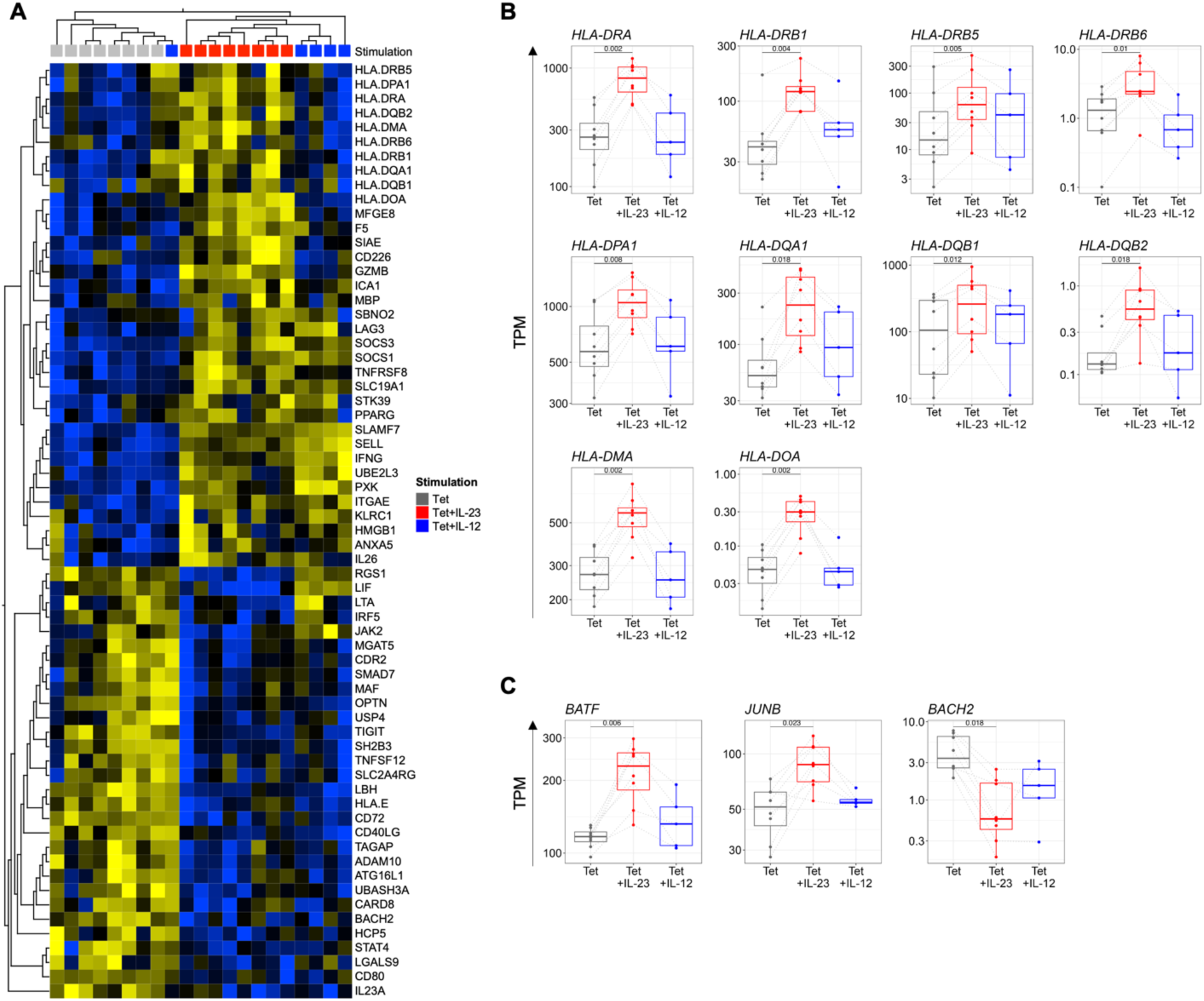
IL-23 regulates MHC class II and AP-1 transcription factor genes in MAIT cells. PBMCs were activated with Tet in the presence or absence of IL-23 or IL-12 for 6 days, CD3^+^CD161^+^Va7.2^+^ MAIT cells were sorted, and RNA-sequencing was performed. (n=5-8). (**A**) Heatmap of the expression of 65 genes associated with autoimmune pathways. Grey: Tet, red: Tet+IL-23, blue: Tet+IL-12. (**B** and **C**) Boxplots of the expression of HLA genes (**B**) and of AP-1 transcription factor genes (**C**). Differential gene expression analysis (limma, adjusted p-values, TPM: transcripts per million)

We also observed that IL-23 affected several members of the AP-1 family of transcription factors. IL-23 enhanced expression of the transcription factors *BATF* and *JUNB* and downregulated *BACH2* expression in MAIT cells (**Figure 5C**). GWAS have shown that variants in *BACH2* are associated with both autoimmune and chronic inflammatory diseases, including SpA, MS, RA, CD and T1D ^43^. BACH2 promotes the differentiation of stem-like CD8^+^ T cells and antagonizes the development of CD8^+^ effector T cells by controling access of AP-1 transcription factors to enhancers and the suppression of effector molecules such as granzymes ^44,45^. Previous studies have demonstrated that Batf^−/−^ and Junb^−/−^ mice are protected from experimental autoimmune encephalomyelitis (EAE), collagen-induced arthritis (CIA) and colitis by inhibiting the differentiation of Th17 cells ^46–49^. Thus, in MAIT cells, IL-23 appears to affect the balance of two opposing transcription factors, BATF and BACH2.

### IL-23 tunes the chromatin landscape in MAIT cells

To elucidate the mechanisms by which IL-23 affects the transcriptional program of MAIT cells, we performed “Assay for Transposase-Accessible Chromatin followed by sequencing” (ATAC-seq) to investigate changes in chromatin accessibility in cells stimulated with Tet in the presence or absence of IL-23. We identified a total of 27,678 accessible chromatin regions, 5% of which (1491) exhibited differential accessibility (DA) upon addition of IL-23. 1032 regions gained accessibility in the presence of IL-23 compared to Tet alone, while 459 showed loss of accessibility, indicating bidirectional effects of IL-23 on chromatin accessibility at the genome-wide level (**Table S9**). Only 29% of the DA regions were located within distal intergenic elements, while the majority (69.2%) were associated with protein-coding genes, being located at promoters (18.3%) or within introns (46.2%) (**Figure S5A**), suggesting that IL-23 signaling can regulate gene expression by affecting chromatin conformation. Gains of chromatin accessibility were found across various regulatory elements, such as promoter regions (e.g. *GZMB or SOCS3*, **Figure 6A**, **B**), intergenic regions (e.g. *IFNG*-*IL26* locus, **Figure 6C**) or introns (*BATF*, **Figure 6D**). Consistent with reduced levels of *BACH2* gene expression in MAIT cells stimulated in the presence of IL-23, we found a loss of chromatin accessibility at the promoter and in an intron of the *BACH2* gene (**Figure 6E**).

**Fig. 6.**
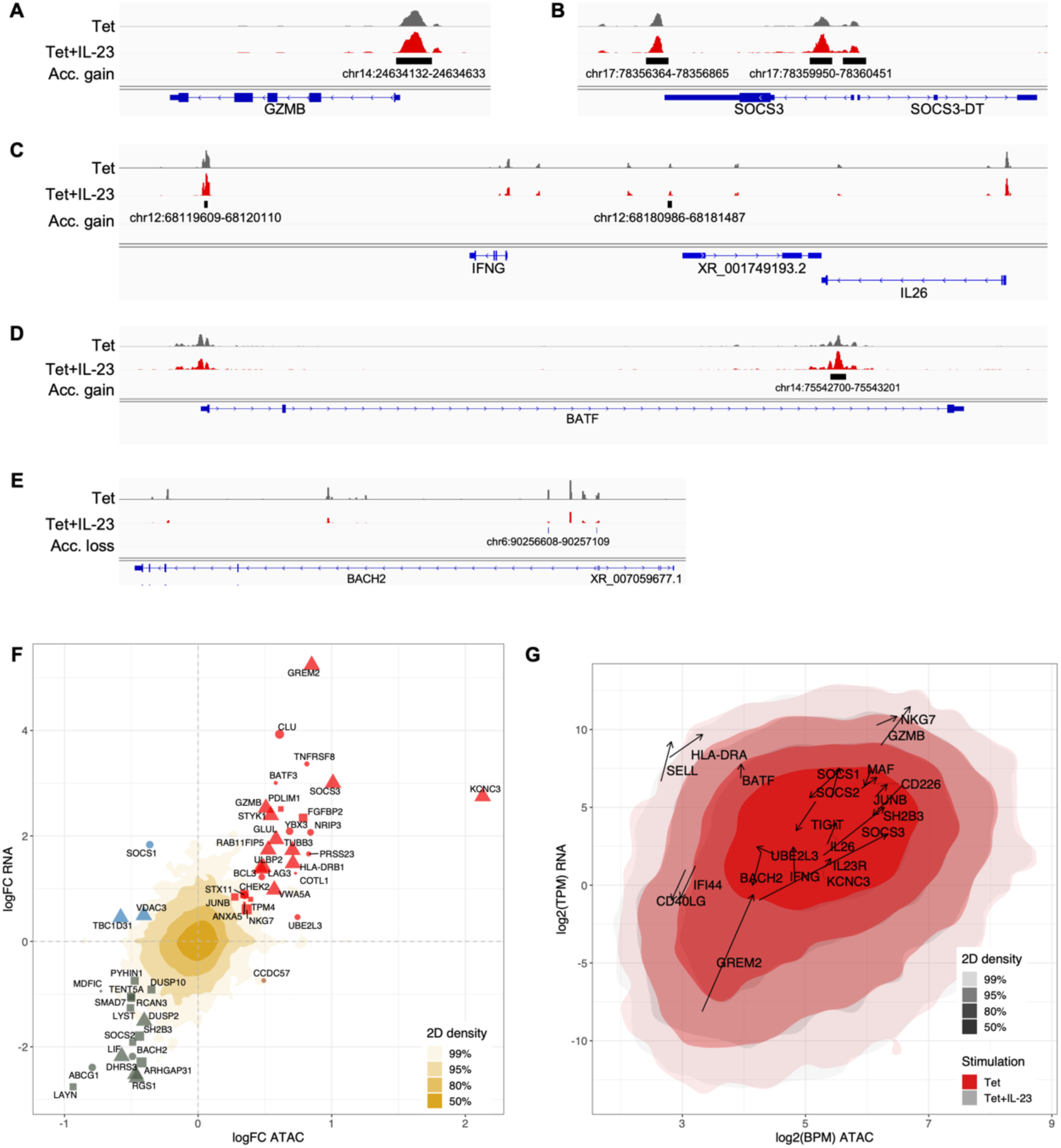
IL-23 regulates the chromatin landscape in MAIT cells. (**A** to **E**) Genome browser tracks of ATAC-seq at indicated loci for the Tet and Tet+IL-23 conditions. One representative replicate is shown for each condition. Significant gains or losses of accessibility are displayed as a separate track. (**F**) Scatter plot of changes in Tet+IL-23 vs. Tet conditions in gene expression (log2-fold change, y axis) as a function of changes in chromatin accessibility (log-fold change, x axis). Each dot corresponds to a differentially expressed and accessible peak-to-gene pair. Red: increase in both accessibility and expression, grey: decrease in both, yellow: increased accessibility, decreased expression, blue: decreased, accessibility, increased expression. Shape indicates position of the DA region relative to the gene’s TSS (triangle: overlapping, square: upstream, circle: downstream). Size is anticorrelated to the distance from the TSS. When several DA regions can be associated to a gene within 10kb, the closest to the TSS is represented. Golden shading depicts the 2D density plot of all log-fold change values. (**G**) 2D density plots of chromatin accessibility (log2BPM) and RNA expression (log2TPM) in Tet and Tet+IL-23 conditions, overlaid. Selected genes are displayed; arrows indicate direction and magnitude of changes in Tet+IL-23 vs. Tet.

Consistent with the functional relationship between the modulation of chromatin accessibility induced by IL-23 and gene expression, there was a significant overlap (105 genes, hypergeometric test, *P* = 4.752784e-19, **Figure S5B, Table S10**) between differentially expressed genes and genes associated with DA regions within 50kb of their transcription start site. Among these genes, transcriptional regulators such as *JUNB*, *BACH2*, *BATF* and *BATF3* and signaling molecules, such as *SOCS2* and *SOCS3,* displayed coordinated chromatin accessibility and gene expression (**Figure 6F**). We found the largest change in chromatin accessibility at the gene encoding the potassium channel *KCNC3* and this gene was also strongly induced by IL-23 (**Table S5**). The genes encoding cytotoxic molecules *GZMB* and *NKG7* also displayed high levels of chromatin accessibility and gene expression in MAIT cells stimulated with Tet alone, which were further increased when cells were stimulated in the presence of IL-23 (**Figure 6G**), providing further evidence for the important role of IL-23 in enhancing cytotoxic functions of MAIT cells.

To explore whether IL-23 treatment may affect transcription factor (TF) binding in tetramer-stimulated MAIT cells, we performed in-silico footprinting to predict TF binding events within accessible chromatin regions ^50^. We identified that stimulation in the presence of IL-23 results, at the genome-wide scale, in increased footprinting of several AP-1 family members, such as JUN, FOS, and BATF and decreased binding of zinc finger TFs, such as KLF12, SP1 and PATZ1, to their respective targets (**Figure 7A**, **Table S11**). Concomitant changes in gene expression and predicted TF binding were observed for BATF, JUNB, and BACH2 (**Figure 7B**), supporting a central role for these AP-1 factors in IL-23 signaling in MAIT cells.

**Fig. 7.**
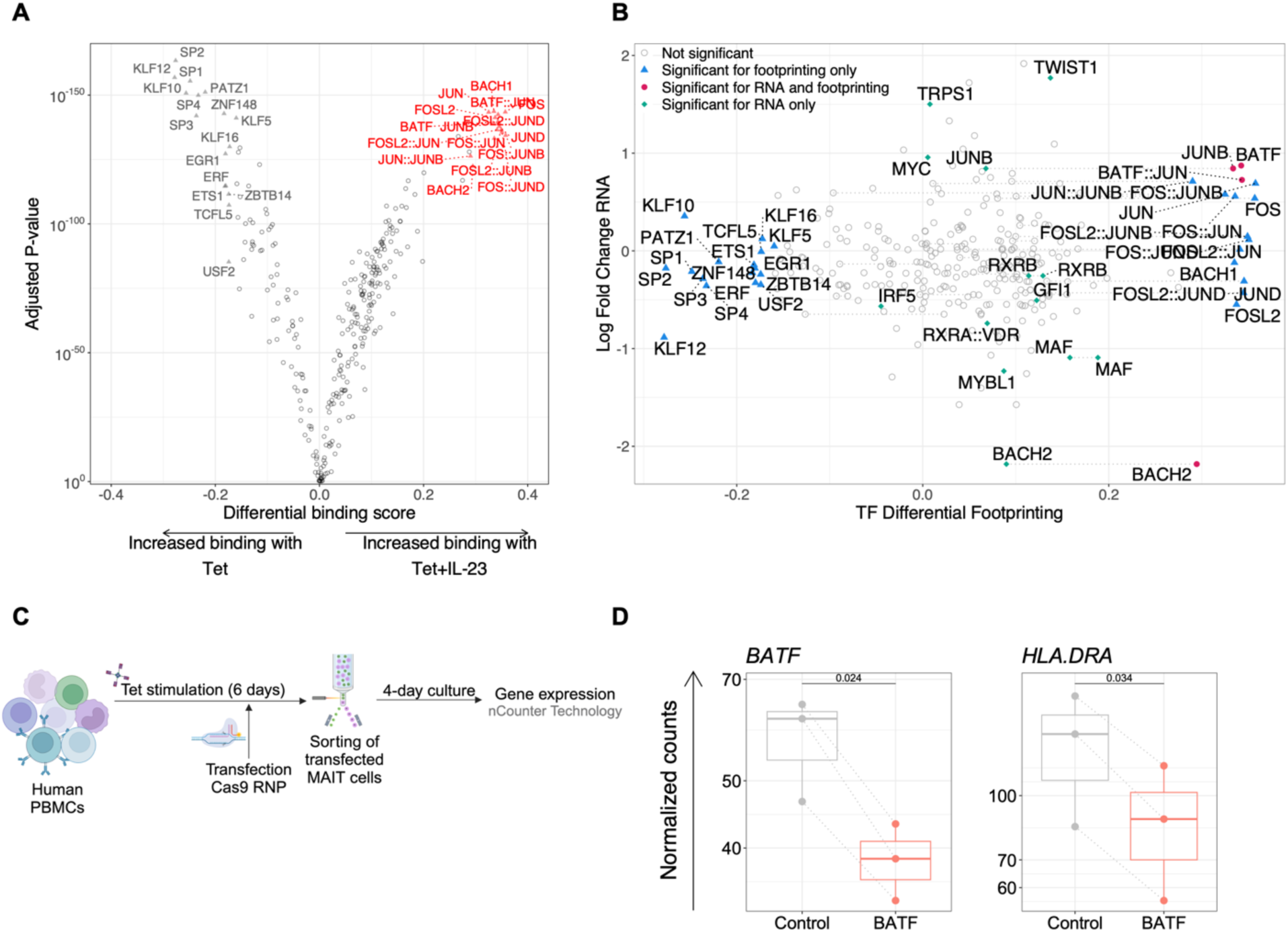
IL-23 signals in MAIT cells are mediated by the AP-1 transcription factors BATF, JUNB, and BACH2 **(A)** Genome-wide *in silico* footprinting analysis was computed on ATAC-seq data from MAIT cells activated with Tet±IL-23. Volcano plot of the genome-wide transcription factor binding changes. Each dot represents one motif, empty dots indicate non-significant events. (**B**) Scatter plot displaying log2-fold change of gene expression against differential binding score for each of the transcription factors used for footprinting. In the case of heterodimers, the gene expression log2-fold change corresponds to the mean of the two genes. Grey dashed lines link the different TF binding prediction for TF binding to several motifs. (**C**-**D**) PBMC were activated with Tet for 6 days. At day 5 of culture, PBMC were transfected with fluorescently labelled Cas9 RNP targeting BATF (Orange) or non-targeting (Control, grey). At day 6, transfected MAIT cells (CD3^+^CD161^+^Va7.2^+^ ATTO550^+^) were sorted, cultured for 4 additional days and gene expression was assessed using nCounter technology (n=3). (**C**) Schematic representation of the experimental design. (**D**) Gene expression (normalized RNA counts) of *BATF* and *HLA-DRA* (p-values, paired t test).

### Control of MAIT cell functions via the transcription factor BATF

We next asked whether BATF may be a key regulator of IL-23 functions in MAIT cells. To suppress BATF expression in MAIT cells, we targeted the BATF gene using CRISPR-Cas9-mediated gene editing (**Figure 7C**). *BATF* expression was reduced in MAIT cells targeted with a BATF gRNA when compared to those targeted with a control (non-targeting) gRNA (**Figure 7D**). BATF targeting resulted in the downregulation of the IL-23 target gene *HLA-DRA* (**Figure 7D**), suggesting that this AP-1 transcription factor constitutes an important regulatory node for the expression of IL-23 target genes.

## Discussion

Most of the knowledge of IL-23 biology has been obtained by the study of IL-23 function in conventional CD4^+^ T cells, in particular in mouse models. Much less is known of IL-23 function in humans, specifically in innate-like T cell populations. In this work we show that in peripheral blood from healthy donors, MAIT cells are the population with the highest frequency of IL-23R-expressing cells. We also detected IL-23R surface expression on a sizable fraction of CD3^+^CD56^+^ and γδ T cells, indicating that innate-like T cells may be important mediators of IL-23 function. These data extend our previous observation that circulating MAIT cells from axial SpA patients expressed higher levels of *IL23R* transcripts than γδ, CD4^+^ and CD8^+^ T cells ^18^.

IL-23 and IL-12 are structurally related cytokines, yet, IL-23p19, but not IL-12p35 knock-out mice were protected from experimental models of autoimmune inflammation in the brain, gut and joints ^51^. These results indicated that only IL-23 but not IL-12 is involved in autoimmunity in mice. Recent results, however, have challenged this notion, showing that IL-12-induced, IL-23R-expressing “Th1-like” cells can cause colitis in a mouse model ^39^, supporting a role for both IL-12 and IL-23 signaling in autoimmune inflammation. We therefore sought to compare the effects of IL-23 and IL-12 stimulation in human MAIT cells. In human and mouse CD4^+^ T cells, IL-23 has been shown to enhance IL-17A production ^11,52^. However IL-23 only modestly increased secretion of IL-17A by stimulated MAIT cells, but had instead substantially stronger effects on the secretion of IFN-γ, IL-10, IL-21 and granzyme B. Of note, IL-12 not only enhanced IFN-γ production but also strongly increased secretion of IL-17A, IL-18 and IL-1β. These findings blur the notion of the canonical roles of IL-12 and IL-23 as regulators, respectively, of type 1 and type 17 immune responses. In addition, it shows that MAIT cells can secrete pro-inflammatory cytokines typically produced by myeloid cells, such as IL-18 and IL-1β. The mechanism by which IL-12 enhances IL-17A secretion by human MAIT cells is currently unclear, but may be mediated by the increased production of IL-18, which has been shown to increase IL-17 production by MAIT cells, in particular in combination with IL-7 ^17^. Our results are consistent with previous data demonstrating that stimulation of human MAIT cells in the presence of IL-23 increases the frequency of IL-17A-producing cells ^53^.

Comparing the effects of IL-23 and IL-12 on tetramer-stimulated MAIT cells revealed that IL-12 affected the expression of considerably more genes (1013) than IL-23 (607). Nevertheless, in MAIT cells activated in the presence of IL-23, we found a clear enrichment of pathways associated with “classical” autoimmune diseases characterized by the presence of autoantibodies, such as MS, SLE and T1D. This was somewhat unexpected because genetic variants in *IL23R* and other genes in the IL-23 pathway are predominantly associated with Crohn’s disease, psoriasis and axial SpA ^10^. These are seronegative diseases characterized by autoinflammatory components and a strong involvement of the innate immune system, rather than B and T cells ^42^. Enrichment of pathways associated with classical autoimmunity can be explained by the fact that IL-23, but not IL-12, upregulates several genes encoding MHC class II molecules. The function of MHC class II molecules on T cells is currently not known, although HLA-DR has been used as a an activation marker of human T cells for many years. Whether MHC class II expressed on T cells may have an antigen-presenting role remains to be demonstrated.

IL-23-driven enhancement of MHC class II gene expression was confirmed by the analysis of stimulated MAIT cells at the single-cell level. We noted that upregulation of class II genes was not restricted to specific clusters of MAIT cells. As a matter of fact, our combined analysis of transcriptomes and epitopes at the single-cell level revealed only a limited heterogeneity of MAIT cells from peripheral blood. In particular, we did not find evidence for the presence of distinct MAIT1 or MAIT17 subsets, which had been identified in the mouse ^25,35,54,55^. Our data are consistent with more recent studies performed with human MAIT cells that did not identify specific MAIT1 or MAIT17 subsets ^29,37,56^.

In addition to regulating MHC class II gene expression, we found that IL-23 regulated several members of the AP-1 family of transcription factors, such as *BATF*, *JUNB*, and *BACH2*. Mouse models have demonstrated the importance of BATF and JUNB for Th17 cell differentiation and function ^46–49^. Targeting BATF in human MAIT cells using CRISPR-Cas9 resulted in a significant reduction of *HLA-DRA* expression, supporting a role for BATF as a transcriptional effector of IL-23 signaling in these cells. Future work will determine further *BATF* target genes in human MAIT cells.

A limitation of our study is that, at the single-cell level, we could reproducibly detect secretion only of granzyme B, perforin and IFN-γ by MAIT cells stimulated with the MR1/5-OP-RU tetramer. Stimulation with phorbol 12-myristate 13-acetate (PMA) and ionomycin allowed the detection of additional secreted molecules, however, these strong stimulation conditions override any effects of IL-23 and IL-12 on MAIT cells (data not shown).

In conclusion, we show that MAIT cells are a key cellular target of IL-23 and of critical importance to mediate the functions of this cytokine in immunity to infections and the pathogenesis of chronic inflammatory diseases.

## Acknowledgments

MR1/5-OP-RU tetramers were generously provided by Dr. Sidonia Eckle and by the NIH tetramer facility. The MR1 tetramer technology was developed jointly by Dr. James McCluskey, Dr. Jamie Rossjohn, and Dr. David Fairlie, and the material was produced by the NIH Tetramer Core Facility as permitted to be distributed by the University of Melbourne.

We thank Valérie Hélin (DARRI, Institut Pasteur) for help in project organization and management. We acknowledge the help of Youssef Ghorbal and the HPC Core Facility of the Institut Pasteur for this work. Sara Bragado Alonso and Catia Cerqueira (EMEA, Bruker Cellular Analysis) assisted with the analysis of IsoPlexis data.

This work was supported by institutional funds from Institut Pasteur, grants from the Fondation de la Recherche Médicale (Equipe FRM, EQU202303016264) and from Janssen Pharmaceuticals (Madeleine project) to LR. LC is supported by a PhD fellowship from the Université Paris Cité and JS-B is supported by a PhD fellowship from the PPU program.

## Author contributions

Conceptualization: LC, TS, EB, LR

Methodology: LC, TS, HY-C, MH, LR

Investigation: LC, TS, HY-C, CL, JM, JS-B, AD

Resources: MR1/5-OP-RU tetramers: HM, SBGE, generation of anti-IL-23R antibody: AF, CG, BJS, RP, DJC

Data analysis: LC, TS, VG, SM, VB, HL-M, DC, LC

Visualization: LC, VG, VB, SM, DC, EB, LR

Data Management: NP

Funding acquisition: EB, LR

Supervision: EB, LR

Writing – original draft: LC, SM, VB, DC, EB, LR

Writing – review & editing:

## STAR★Methods

### Resource availability

#### Lead contact

Further information and requests for resources and reagents should be directed to and will be fulfilled by the lead contact, Lars Rogge.

#### Materials availability

Requests for the anti-IL-23R monoclonal antibody used in this study should be sent to Carrie Greving.

#### Data and code availability

- Raw sequencing data for RNA-seq, ATAC-seq and CITE-seq will be available on Gene Expression Omnibus.
- Generation and characteristics of the anti-IL-23R monoclonal antibody used in this study are described in a separate manuscript, in preparation and available on request from Dr. Carrie Greving.
- This paper does not report original code.
- Any additional information required to reanalyze the data reported in this work paper is available from the lead contact upon request.

#### Experimental model and study participant details

Blood samples from healthy donors were obtained from Etablissement Français du Sang (EFS, Paris, France)

#### Method details

##### Assessment of cell surface IL-23R expression by spectral flow cytometry

Peripheral blood mononuclear cells (PBMC) were freshly isolated by density gradient centrifugation and prepared as single cell suspension in staining buffer (phosphate-buffered saline (PBS), 1% fetal bovine serum (FBS), 5% bovine serum albumin (BSA)). Cells were stained with Live/Dead Fixable Blue (1:100) and Fc receptor blocker for 5 min on ice. Cells were stained with either anti-IL-23R monoclonal antibody or IgG2a isotype control for 1 h on ice. After washing, cells were incubated with biotinylated anti-mouse IgG2a for 30 min on ice. Cells were then washed and incubated with streptavidin-PE for 15 min on ice. After washing, cells were stained with cell surface marker antibodies (**Table S1**) for 30 min at room temperature (RT). Then, after washing, cells were fixed with 4% paraformaldehyde for 15min at RT. Cells were acquired on a Sony ID7000. Unmixing was performed using the Sony ID7000 Software, and analysis was performed using FlowJo as described ^57^.

##### MAIT cell stimulation

PBMC were isolated by density gradient centrifugation and cultured in RPMI supplemented with 10% FBS, 1% penicillin/streptomycin and 20UI/mL of IL-2. MR1/5-OP-RU tetramers (unlabeled) were added the first day of culture at 40pM, and cells were cultured for 6 days. When indicated, IL-23 or IL-12 were added at the concentration of 20 ng/mL.

##### FACS sorting of MAIT cells

After stimulation, cells were prepared as single cell suspensions in staining buffer (PBS, 1% FBS) and stained for 20 min at 4°C with eF520 fixable viability dye (1:100) and surface marker antibodies: anti-CD3-BV711 (1:20), anti-TCRVa7.2-APC (1:50), and anti-CD161-BV421 (1:50) or MR1/5-OP-RU-BV421 tetramers (1:600). MAIT cells were sorted, as indicated, as CD3^+^TCRVa7.2^+^CD161^high^ or CD3^+^MR1/5-OP-RU^+^ on a BD FACS Aria III.

##### Assessment of cytokine secretion

Secretion in the supernatant of 20 proteins (**Table S2**, ProcartaPlex custom kit) was measured following manufacturer’s instructions using the DropArray technology (Curiox). Briefly, the DropArray plate was blocked with PBS-1% BSA and washed. Samples and standards were incubated with capture beads for 2h at RT. After washing, detection antibody mix was added and incubated for 30min at RT. After washing, streptavidin-PE was added and incubated for 30min at RT. Samples were acquired on a Bioplex200. Analysis was performed using the Bio-Plex Manager Software. Concentration of each analyte was calculated using a 5-PL regression curve. Individual out of range values were replaced with the lowest value measured in the data set divided by 2 (lower out of range), or the highest value multiplied by two (upper out of range). Before plotting, the data was centered and scaled (row Z-scores), the rows and columns reordered according to a hierarchical clustering analysis based on the Euclidean distance with Ward agglomerative method. Adjusted p-values were obtained by nonparametric pairwise Wilcoxon rank-sum test (between each pairs of groups), corrected wih Benjamini-Hochberg multiple testing corrections applied on all the tests performed (on all cytokines).

##### Single-cell secretome profiling

MAIT cells were activated in PBMC cultures for 6 days in the presence or absence of IL-23 or IL-12. At day 6, MAIT cells (CD3^+^MR1/5-OP-RU^+^) were sorted and rested overnight in RPMI. MAIT cells were re-stimulated for 1h with MR1/5-OP-RU tetramers in the presence or absence of IL-23 or IL-12. Cells were then washed, incubated with membrane stain (Isoplexis) for 15min at 37°C. After washing, cells were resuspended in RPMI with or without IL-23 or IL-12 at a concentration of 10^6^cells/mL. 30µL of the cell suspensions were then loaded on IsoCode chips. Data were acquired on an Isolight instrument, and analysed using the IsoSpeak Software.

##### RNA-sequencing

After 6 days of activation in PBMC cultures, MAIT cells (CD3^+^ TCRVa7.2^+^ CD161^high^) were sorted and RNA was isolated using RNeasy Mini or Micro Kit. mRNA libraries were generated and sequenced at Novogene. Libraries were sequenced on the Illumina NovaSeq 6000 (150bp, paired-end, 20M reads per sample). Reads were mapped to the reference human genome (hg38) using STAR. FeatureCounts was used to estimate gene expression for each sample. All samples passed the quality control which was performed on raw reads using FastQC.

Differential analysis was performed using limma on variance stabilized data ^58^, with an additional step to estimate the between donor correlation. Reported p-values were adjusted with Benjamini-Hochberg correction for multiple testing. Functional analysis (GSEA) was performed using WebGestaltR on disease related databases ^59^.

##### ATAC-sequencing

After 6 days of activation in PBMC cultures, MAIT cells (CD3^+^ TCRVa7.2^+^ CD161^high^) were sorted and ATAC-seq was performed on 90000 cells. Libraries were prepared using the ATAC-seq kit from Active motif according to manufacturer’s instructions. Briefly, nuclei were isolated by adding 100μL ice cold ATAC-lysis buffer to the cell pellet. After centrifugation (2800 rpm, 10 min 4°C), nuclei were incubated with the tagmentation master mix in a shaking heat block for 30min at 37°C with agitation at 800rpm. DNA was then purified and amplified for 10 cycles using indexed primers, and size-selected using SPRI bead solution. Quantification of the libraries was done using the Qubit dsDNA HS Assay Kit. Quality control of the size distribution of the PCR enriched library fragments was performed using D5000 DNA chip (Agilent Technologies). The libraries were sequenced on the Illumina NovaSeq 6000 (150bp, paired-end, 80M reads per sample).

Read quality was checked with FastQC. Raw sequencing reads were trimmed using Cutadapt [https://doi.org/10.14806/ej.17.1.200] to remove sequencing adapters and low-quality bases (Q<30) and reads shorter than 25bp after trimming were discarded. Cutadapt was further used to crop the remaining reads down to 50bp. Reads were aligned to human genome hg38 in paired-end mode using Bowtie2 with the following parameters: *--very-sensitive -X 2000 --dovetail --no-mixed --no-discordant.* Duplicates were removed using Picard MarkDuplicates, as well as reads mapping to mitochondrial chromosome and to blacklisted regions as defined in ^60^. Following bam to bed conversion using Bedtools, peaks were called separately for each sample using MACS2 with the following parameters: *--nomodel --shift -100 --extsize 200 --keep-dup all -g hs --call-summits*. BigWig tracks for IGV visualization were produced using deepTools, using BPM normalization over 10bp bins.

To produce a consensus peak list, peaks were combined using the iterative peak merging approach as described in ^61^. This consensus peak list was used for both differential accessibility and in silico transcription factor footprinting analyses.

Differential accessibility was performed using DESeq2, regressing out the donor effect. False Discovery Rate (FDR) <0.05 was used to define significant differential accessibility. BPM counts were computed over 501nt long bins. Peak annotation was performed using rGREAT under the basal+extension rule (5+1kb, 1M extension), using RefSeq Select TSS annotation. The analysis was performed using R 4.3.

##### In silico transcription factor footprinting

TOBIAS was run with default settings on merged biological replicates, using as for motif database a merge of JASPAR (JASPAR 2022 release, 727 TFs motifs) and selected motifs from HOCOMOCO (HOCOMOCOv11_core_HUMAN_mono_jaspar_format.txt) ^50^. TOBIAS predicted footprints were further filtered to only retain the transcription factors for which genes were expressed in the biological conditions under scrutiny based on the RNA-seq data (threshold for expression: mean TPM across the donors = 1). The analysis was performed using R 4.3.

##### CITE-seq

MAIT cells were activated in PBMC cultures for 6 days in the presence or absence of IL-23 or IL-12. At day 6, 10^6^ cells were incubated with Total-seq C antibodies (Biolegend) and surface fluorochrome-conjugated antibodies for MAIT cell sorting for 30min at 4°C. MAIT cells were sorted as CD3^+^MR1/5-OP-RU^+^ and loaded onto a chromium 5’ chip following the Single Cell 5’ Kit and run on the 10X Chromium. Gene expression (GEX), V(D)J and cell surface protein libraries preparation and acquisition of sequencing reads were generated according to the manufacturer’s recommendations using the User Guide CG000330 Rev D, Chromium Next GEM Single Cell 5’ Reagent Kits v2. The libraries were sequenced on the Illumina NovaSeq 6000 (150 bp, paired-end).

The raw sequencing data, comprising antibody capture, TCR, and gene expression, was provided as sample-demultiplexed fastq files. The generation of gene expression and antibody capture count matrices, as well as TCR VDJ Tables, was performed using cellranger multi software with refdata-gex-GRCh38-2020-A, refdata-cellranger-vdj-GRCh38-alts-ensembl-7.0.0., and 10XTotalSeq as references for the GEX, the V(D)J, and the CITE-seq data, respectively.

The analysis was performed in R (v4.2.3) with the Seurat environment. First, for each sample, the three modalities were integrated into the same Seurat object. Gene expression and antibody matrices were considered as different Assays, while for the TCR sequences, scRepertoire was used to read the V(D)J Table (filtered_config_annotations.csv) prior to integration into the meta.data of the Seurat object.

The cell filtering process involved basic metrics, such as the number of genes (>750), unique molecular identifiers (UMI>1000), and the maximum percentage of mitochondrial gene expression (<10%). Additional criteria for cell filtering included the expression of selected genes, such as positive expression of CD3 genes and TRAV1-2 genes at the gene expression (gex) level, antibody capture, as well as for TRAV1-2 at VDJ levels.

Each sample was processed individually according to the following standard workflow: Normalization using SCTransform with 3000 variable features and the percentage of mitochondrial expression regressed out; Linear dimension reduction using PCA (with mitochondrial, ribosomal and TCR genes removed from the highly variable genes), and non-linear dimension reduction using UMAP (with the first 30 PCs); Graph-based clustering and the Louvain algorithm was used to find communities at different resolutions.

Once processed, individual samples were merged together using the harmony integration method and grouped by sample id. Harmony was applied separately to gene expression and antibody capture data. The corresponding corrected matrices were then used as input in the weighted nearest neighbors (WNN) algorithm. This weighted neighbors graph was used as input for the non-linear reduction in the RunUMAP Seurat function and for the graph-based clustering with the Smart Local Moving (SLM) algorithm option in the FindCluster Seurat function.

Signature scores (Cytotoxicity, IFN, MAIT1, MAIT17, Th1 and Th17) were computed using UCell library (v2.4.0).

##### Mowgli Integration of CITE-seq data and identification of specific factors

The RNA and antibody modalities of the CITE-seq data were integrated with Mowgli ^41^. We performed one integration per donor. RNA counts were log-normalized, while centered-log ratio (CLR) normalization was used for antibodies. Only the top 1500 highly variable genes were used in the integration, while all the 137 high-quality antibodies were kept. The standard parameters for Mowgli Integration of CITE-Seq data were applied, except for the number of latent dimensions. This number was set to 100 to increase the granularity of the biological signal between cells treated with Tet and cells treated with Tet+IL-23.

For the Tet+IL-23 specific latent factors associated with each donor, Spearman correlation was performed on the matrix elements identified by Mowgli notation. For RNA, only genes in common between Donor 1 and Donor 2 or 3 were selected. For ADT, all antibodies used for each Mowgli Integration were selected.

##### CRISPR-Cas9 editing

crRNAs, tracrRNA-ATTO550, Cas9 and electroporation enhancer were obtained from IDT. crRNA was designed using the CHOPCHOP online tool ^62^ and the IDT design checker and chosen for highest target specificity and lowest number of off-targets.

Guide RNAs (gRNAs) were formed by heating 2µL of 200µM-crRNA, 2µL of 200uM-tracrRNA and 5µL of Duplex Buffer at 95°C for 5min, and allowed to cool down at RT. Ribonucleoproteins (RNP) were then formed by incubating 8µL of gRNA with 5µL Cas9 nuclease (36uM) for 20min at RT. 10^7^ PBMCs activated for 5 days with MR1/5-OP-RU tetramers were harvested and resuspended in 80µL of T buffer (Invitrogen), with 13µL RNP and 20µL electroporation enhancer (10.8µM). Electroporation was performed with the Neon Electroporation system, with 3 pulses of 10ms at 1800V. 24h after electroporation, transfected MAIT cells (TCRVα7.2^+^CD161^high^ATTO550^+^) were sorted and cultured for 4 days in RPMI supplemented with 10% FBS, 1% penicillin/streptomycin and 20UI/mL of IL-2. RNA was isolated using RNeasy Micro Kit.

##### Gene expression analysis with nCounter

RNA concentration was measured using Qubit RNA HS Assay Kit. 50ng of total RNA from each sample were analyzed using Nanostring nCounter technology with Immunology V2 panel and a custom Plus panel as described ^63^. Statistical analysis (paired t test) and graphical representation were performed in R (v4.2.2).

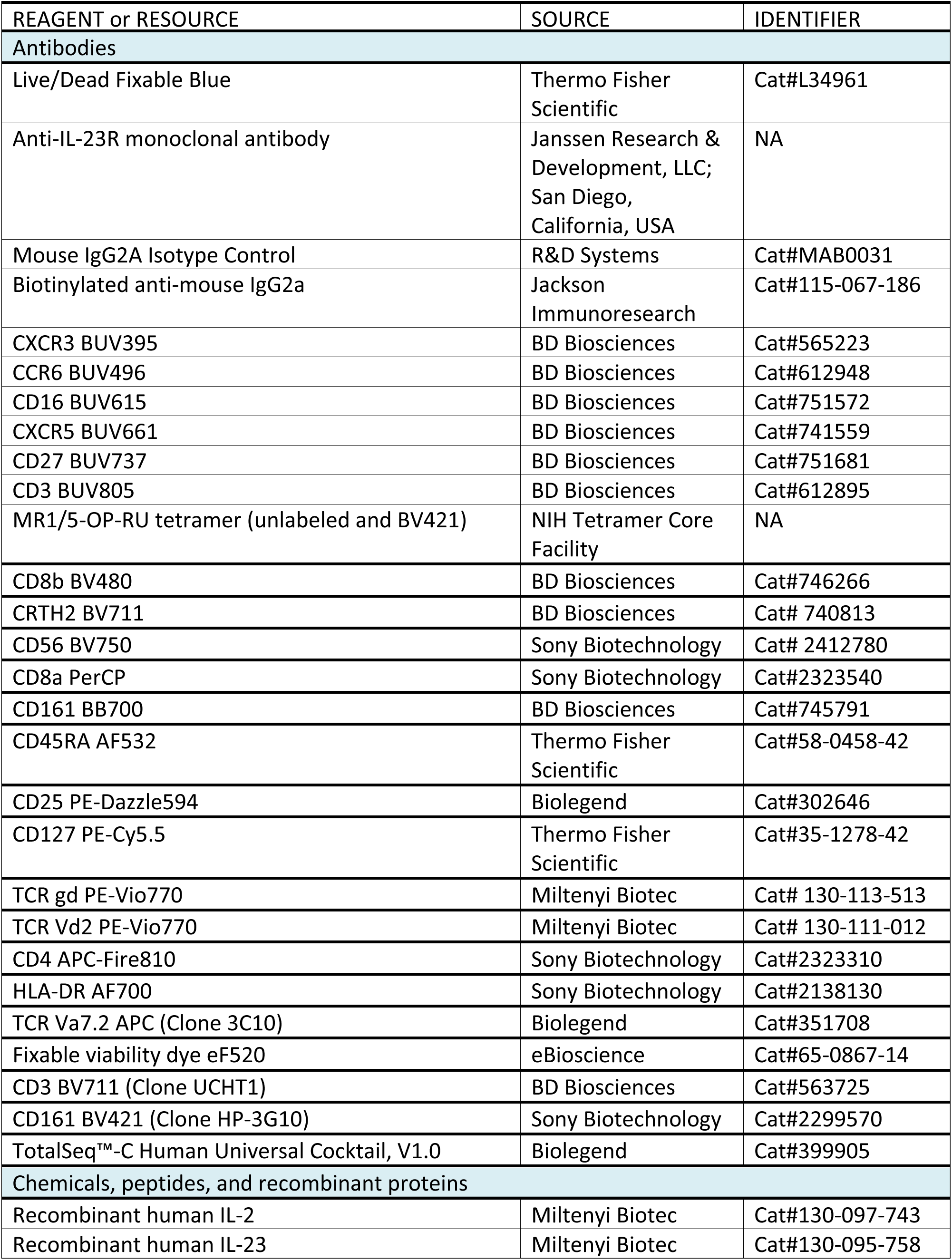
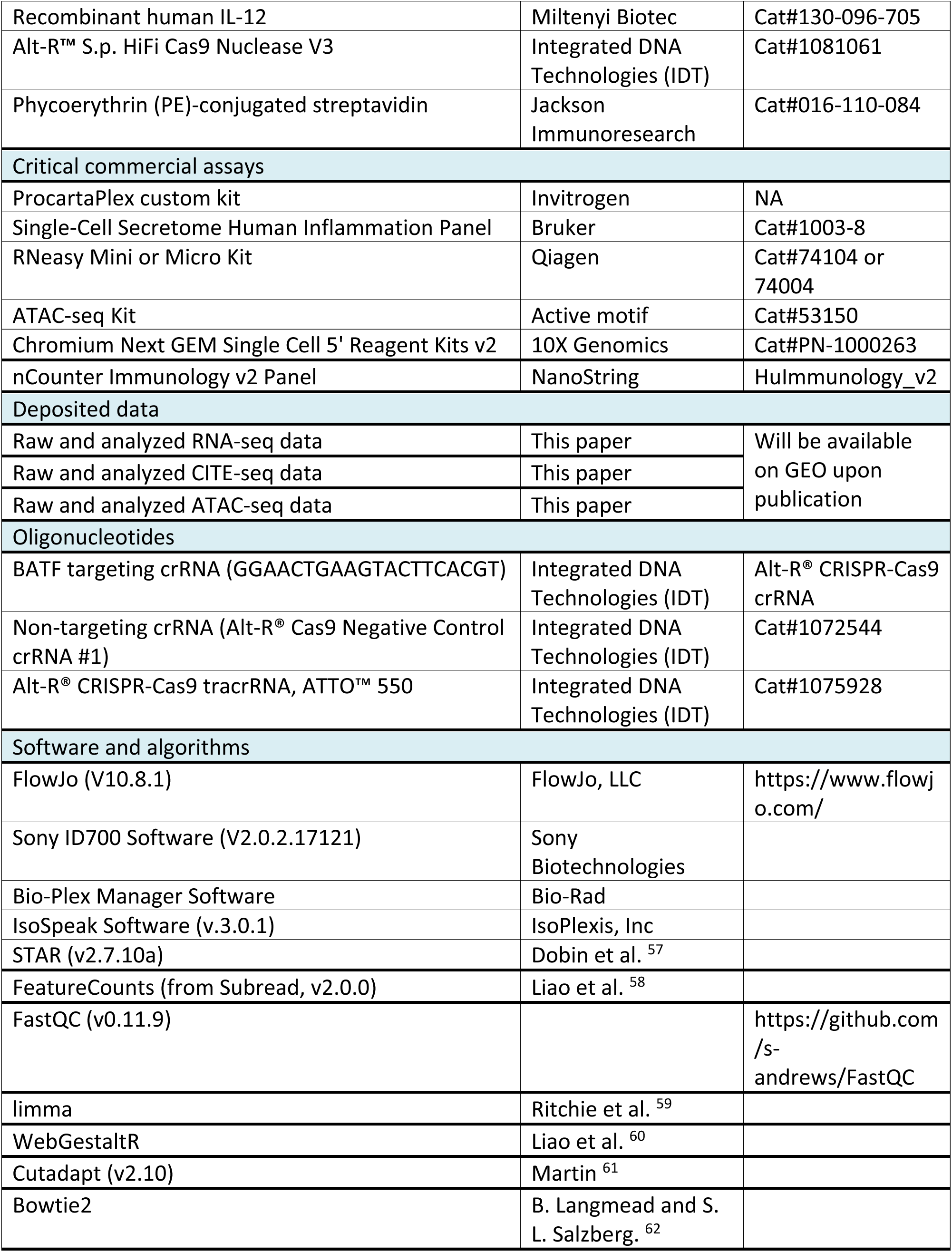
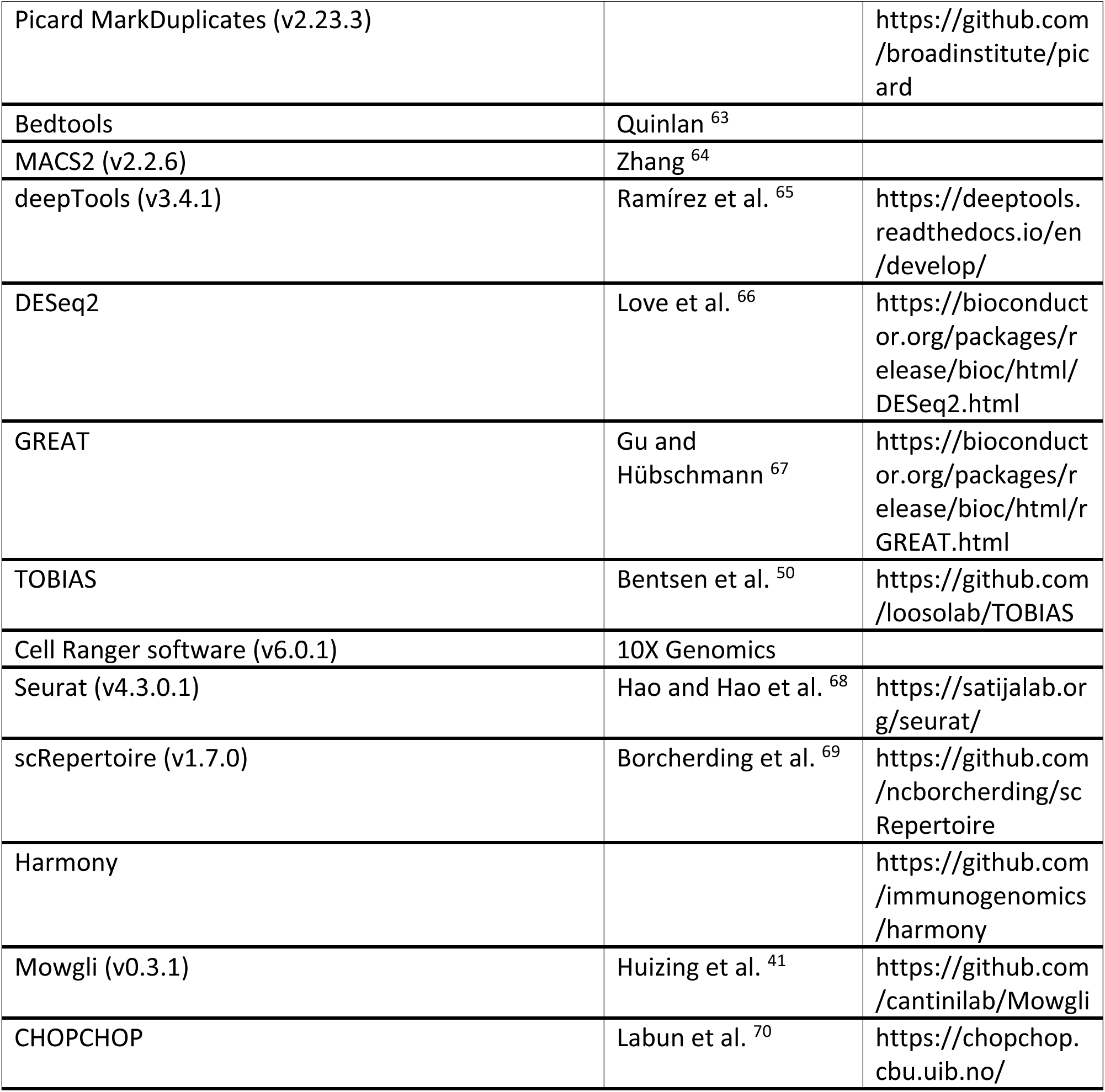
Key resources table

## Supplementary Materials

### Supplementary figures

Figure S1: CITE-seq analysis, related to Figure 2

Figure S2: CITE-seq analysis, related to Figure 2

Figure S3: CITE-seq analysis, related to Figure 2

Figure S4: CITE-seq analysis, related to Figure 2

Figure S5: ATAC-seq analysis, related to Figure 6

## Supplementary Tables

**Table S1: Antibody panel for analysis of IL-23R expression on immune cells**

**Table S2: Proteins secreted by MAIT cells, related to** Figure 1. Measurements of cytokines using Luminex technology.

**Table S3: CITE-seq analysis, genes defining clusters, related to Figure 2**

**Table S4: CITE-seq analysis, cell surface markers defining clusters, related to Figure 2**

**Table S5: 607 genes regulated by IL-23 (bulk RNA seq), related to** Figure 3. Limma analysis of differentially expressed genes in Tetramer + IL-23 stimulated cells, relative to stimulation with Tetramer alone (adj Pval ≤0.05)

**Table S6: 1013 genes regulated by IL-12 (bulk RNA seq), related to** Figure 3. Limma analysis of differentially expressed genes in Tetramer + IL-12 stimulated cells, relative to stimulation with Tetramer alone (adj Pval ≤0.05)

**Table S7: Top 20 pathways regulated by IL-23, related to** Figure 3. Results of the over-representation analysis performed on the 607 genes regulated by IL-23 using the GLAD4U database.

**Table S8: Top 20 pathways regulated by IL-12, related to** Figure 3. Results of the over-representation analysis performed on the 1013 genes regulated by IL-12 using the GLAD4U database.

**Table S9: Differential accessibility upon addition of IL-23 (989 gains, 424 losses), related to** Figure 5. This Table reports the chromatin regions identified as differentially accessible comparing cells stimulated in the presence or in the absence of IL-23. Each row corresponds to a DA region. For each region, log2FC, pvalue and adjusted pvalue (pajd) are provided, as well as distance to TSS and gene annotation (basal+extension model). “GAIN” and “LOSS” denotes whether this region exhibits a gain or a loss in accessibility in Tetramer+IL23 stimulated cells, respectively.

**Table S10: Table corresponding to Venn diagram; correspondence of differentially expressed genes and accessible genes, related to Figure S4.** This Table reports the differentially expressed genes (DE) associated with differentially accessible regions (DA) within 50kb of their TSS. Each row corresponds to one DE gene/ DA region association. If several DA regions can be annotated to the same given gene, these are reported as distinct rows.

**Table S11: Genome-wide differential binding predictions for transcription factors, related to** Figure 6. This Table reports, for each TF binding motif under study, its genome wide metrics (differential binding score, adjusted pvalue). Positive differential binding scores indicate higher binding in the Tet+ IL-23 condition, negative scores higher binding in the Tet condition.

## References

1. Lubberts, E. (2015). The IL-23-IL-17 axis in inflammatory arthritis. Nature reviews. Rheumatology 11, 562. 10.1038/nrrheum.2015.128.

2. Philippot, Q., Ogishi, M., Bohlen, J., Puchan, J., Arias, A.A., Nguyen, T., Martin-Fernandez, M., Conil, C., Rinchai, D., Momenilandi, M., et al. (2023). Human IL-23 is essential for IFN-gamma-dependent immunity to mycobacteria. Sci Immunol 8, eabq5204. 10.1126/sciimmunol.abq5204.

3. Krueger, J.G., Eyerich, K., Kuchroo, V.K., Ritchlin, C.T., Abreu, M.T., Elloso, M.M., Fourie, A., Fakharzadeh, S., Sherlock, J.P., Yang, Y.W., et al. (2024). IL-23 past, present, and future: a roadmap to advancing IL-23 science and therapy. Front Immunol 15, 1331217. 10.3389/fimmu.2024.1331217.

4. Cua, D.J., Sherlock, J., Chen, Y., Murphy, C.A., Joyce, B., Seymour, B., Lucian, L., To, W., Kwan, S., Churakova, T., et al. (2003). Interleukin-23 rather than interleukin-12 is the critical cytokine for autoimmune inflammation of the brain. Nature 421, 744–748.

5. Hue, S., Ahern, P., Buonocore, S., Kullberg, M.C., Cua, D.J., McKenzie, B.S., Powrie, F., and Maloy, K.J. (2006). Interleukin-23 drives innate and T cell-mediated intestinal inflammation. J Exp Med 203, 2473–2483. 10.1084/jem.20061099.

6. Murphy, C.A., Langrish, C.L., Chen, Y., Blumenschein, W., McClanahan, T., Kastelein, R.A., Sedgwick, J.D., and Cua, D.J. (2003). Divergent pro- and antiinflammatory roles for IL-23 and IL-12 in joint autoimmune inflammation. J Exp Med 198, 1951–1957. 10.1084/jem.20030896.

7. Yen, D., Cheung, J., Scheerens, H., Poulet, F., McClanahan, T., McKenzie, B., Kleinschek, M.A., Owyang, A., Mattson, J., Blumenschein, W., et al. (2006). IL-23 is essential for T cell-mediated colitis and promotes inflammation via IL-17 and IL-6. J Clin Invest 116, 1310–1316. 10.1172/JCI21404.

8. McGeachy, M.J., Chen, Y., Tato, C.M., Laurence, A., Joyce-Shaikh, B., Blumenschein, W.M., McClanahan, T.K., O’Shea, J.J., and Cua, D.J. (2009). The interleukin 23 receptor is essential for the terminal differentiation of interleukin 17-producing effector T helper cells in vivo. Nat Immunol 10, 314–324.

9. Duerr, R.H., Taylor, K.D., Brant, S.R., Rioux, J.D., Silverberg, M.S., Daly, M.J., Steinhart, A.H., Abraham, C., Regueiro, M., Griffiths, A., et al. (2006). A genome-wide association study identifies IL23R as an inflammatory bowel disease gene. Science 314, 1461–1463.

10. Ellinghaus, D., Jostins, L., Spain, S.L., Cortes, A., Bethune, J., Han, B., Park, Y.R., Raychaudhuri, S., Pouget, J.G., Hubenthal, M., et al. (2016). Analysis of five chronic inflammatory diseases identifies 27 new associations and highlights disease-specific patterns at shared loci. Nat Genet 48, 510–518. 10.1038/ng.3528.

11. Coffre, M., Roumier, M., Rybczynska, M., Sechet, E., Law, H.K., Gossec, L., Dougados, M., Bianchi, E., and Rogge, L. (2013). Combinatorial control of Th17 and Th1 cell functions by genetic variations in genes associated with the interleukin-23 signaling pathway in spondyloarthritis. Arthritis and rheumatism 65, 1510–1521. 10.1002/art.37936.

12. Bianchi, E., and Rogge, L. (2019). The IL-23/IL-17 pathway in human chronic inflammatory diseases-new insight from genetics and targeted therapies. Genes Immun 20, 415–425. 10.1038/s41435-019-0067-y.

13. Langrish, C.L., Chen, Y., Blumenschein, W.M., Mattson, J., Basham, B., Sedgwick, J.D., McClanahan, T., Kastelein, R.A., and Cua, D.J. (2005). IL-23 drives a pathogenic T cell population that induces autoimmune inflammation. J Exp Med 201, 233–240.

14. Sherlock, J.P., Joyce-Shaikh, B., Turner, S.P., Chao, C.C., Sathe, M., Grein, J., Gorman, D.M., Bowman, E.P., McClanahan, T.K., Yearley, J.H., et al. (2012). IL-23 induces spondyloarthropathy by acting on ROR-gammat(+) CD3(+)CD4(-)CD8(-) entheseal resident T cells. Nature medicine 18, 1069–1076.

15. Kenna, T.J., Davidson, S.I., Duan, R., Bradbury, L.A., McFarlane, J., Smith, M., Weedon, H., Street, S., Thomas, R., Thomas, G.P., and Brown, M.A. (2012). Enrichment of circulating interleukin-17-secreting interleukin-23 receptor-positive gamma/delta T cells in patients with active ankylosing spondylitis. Arthritis and rheumatism 64, 1420–1429. 10.1002/art.33507.

16. Venken, K., Jacques, P., Mortier, C., Labadia, M.E., Decruy, T., Coudenys, J., Hoyt, K., Wayne, A.L., Hughes, R., Turner, M., et al. (2019). RORgammat inhibition selectively targets IL-17 producing iNKT and gammadelta-T cells enriched in Spondyloarthritis patients. Nat Commun 10, 9. 10.1038/s41467-018-07911-6.

17. Gracey, E., Qaiyum, Z., Almaghlouth, I., Lawson, D., Karki, S., Avvaru, N., Zhang, Z., Yao, Y., Ranganathan, V., Baglaenko, Y., and Inman, R.D. (2016). IL-7 primes IL-17 in mucosal-associated invariant T (MAIT) cells, which contribute to the Th17-axis in ankylosing spondylitis. Ann Rheum Dis 75, 2124–2132. 10.1136/annrheumdis-2015-208902.

18. Rosine, N., Rowe, H., Koturan, S., Yahia-Cherbal, H., Leloup, C., Watad, A., Berenbaum, F., Sellam, J., Dougados, M., Aimanianda, V., et al. (2022). Characterization of Blood Mucosal-Associated Invariant T Cells in Patients With Axial Spondyloarthritis and of Resident Mucosal-Associated Invariant T Cells From the Axial Entheses of Non-Axial Spondyloarthritis Control Patients. Arthritis Rheumatol 74, 1786–1795. 10.1002/art.42090.

19. Godfrey, D.I., Koay, H.F., McCluskey, J., and Gherardin, N.A. (2019). The biology and functional importance of MAIT cells. Nat Immunol 20, 1110–1128. 10.1038/s41590-019-0444-8.

20. Tilloy, F., Treiner, E., Park, S.H., Garcia, C., Lemonnier, F., de la Salle, H., Bendelac, A., Bonneville, M., and Lantz, O. (1999). An invariant T cell receptor alpha chain defines a novel TAP-independent major histocompatibility complex class Ib-restricted alpha/beta T cell subpopulation in mammals. J Exp Med 189, 1907–1921. 10.1084/jem.189.12.1907.

21. Provine, N.M., and Klenerman, P. (2020). MAIT Cells in Health and Disease. Annu Rev Immunol 38, 203–228. 10.1146/annurev-immunol-080719-015428.

22. Kjer-Nielsen, L., Patel, O., Corbett, A.J., Le Nours, J., Meehan, B., Liu, L., Bhati, M., Chen, Z., Kostenko, L., Reantragoon, R., et al. (2012). MR1 presents microbial vitamin B metabolites to MAIT cells. Nature 491, 717–723. 10.1038/nature11605.

23. Treiner, E., Duban, L., Bahram, S., Radosavljevic, M., Wanner, V., Tilloy, F., Affaticati, P., Gilfillan, S., and Lantz, O. (2003). Selection of evolutionarily conserved mucosal-associated invariant T cells by MR1. Nature 422, 164–169. 10.1038/nature01433.

24. Le Bourhis, L., Martin, E., Peguillet, I., Guihot, A., Froux, N., Core, M., Levy, E., Dusseaux, M., Meyssonnier, V., Premel, V., et al. (2010). Antimicrobial activity of mucosal-associated invariant T cells. Nat Immunol 11, 701–708. 10.1038/ni.1890.

25. Salou, M., Legoux, F., Gilet, J., Darbois, A., du Halgouet, A., Alonso, R., Richer, W., Goubet, A.G., Daviaud, C., Menger, L., et al. (2019). A common transcriptomic program acquired in the thymus defines tissue residency of MAIT and NKT subsets. J Exp Med 216, 133–151. 10.1084/jem.20181483.

26. Constantinides, M.G., Link, V.M., Tamoutounour, S., Wong, A.C., Perez-Chaparro, P.J., Han, S.J., Chen, Y.E., Li, K., Farhat, S., Weckel, A., et al. (2019). MAIT cells are imprinted by the microbiota in early life and promote tissue repair. Science 366. 10.1126/science.aax6624.

27. Provine, N.M., Amini, A., Garner, L.C., Spencer, A.J., Dold, C., Hutchings, C., Silva Reyes, L., FitzPatrick, M.E.B., Chinnakannan, S., Oguti, B., et al. (2021). MAIT cell activation augments adenovirus vector vaccine immunogenicity. Science 371, 521–526. 10.1126/science.aax8819.

28. du Halgouet, A., Darbois, A., Alkobtawi, M., Mestdagh, M., Alphonse, A., Premel, V., Yvorra, T., Colombeau, L., Rodriguez, R., Zaiss, D., et al. (2023). Role of MR1-driven signals and amphiregulin on the recruitment and repair function of MAIT cells during skin wound healing. Immunity 56, 78–92 e76. 10.1016/j.immuni.2022.12.004.

29. Garner, L.C., Amini, A., FitzPatrick, M.E.B., Lett, M.J., Hess, G.F., Filipowicz Sinnreich, M., Provine, N.M., and Klenerman, P. (2023). Single-cell analysis of human MAIT cell transcriptional, functional and clonal diversity. Nat Immunol 24, 1565–1578. 10.1038/s41590-023-01575-1.

30. Corbett, A.J., Eckle, S.B., Birkinshaw, R.W., Liu, L., Patel, O., Mahony, J., Chen, Z., Reantragoon, R., Meehan, B., Cao, H., et al. (2014). T-cell activation by transitory neo-antigens derived from distinct microbial pathways. Nature 509, 361–365. 10.1038/nature13160.

31. Stoeckius, M., Hafemeister, C., Stephenson, W., Houck-Loomis, B., Chattopadhyay, P.K., Swerdlow, H., Satija, R., and Smibert, P. (2017). Simultaneous epitope and transcriptome measurement in single cells. Nature methods 14, 865–868. 10.1038/nmeth.4380.

32. Lanier, L.L., Corliss, B., Wu, J., and Phillips, J.H. (1998). Association of DAP12 with activating CD94/NKG2C NK cell receptors. Immunity 8, 693–701. 10.1016/s1074-7613(00)80574-9.

33. Lanier, L.L., Corliss, B.C., Wu, J., Leong, C., and Phillips, J.H. (1998). Immunoreceptor DAP12 bearing a tyrosine-based activation motif is involved in activating NK cells. Nature 391, 703–707. 10.1038/35642.

34. Koay, H.F., Su, S., Amann-Zalcenstein, D., Daley, S.R., Comerford, I., Miosge, L., Whyte, C.E., Konstantinov, I.E., d’Udekem, Y., Baldwin, T., et al. (2019). A divergent transcriptional landscape underpins the development and functional branching of MAIT cells. Sci Immunol 4. 10.1126/sciimmunol.aay6039.

35. Legoux, F., Gilet, J., Procopio, E., Echasserieau, K., Bernardeau, K., and Lantz, O. (2019). Molecular mechanisms of lineage decisions in metabolite-specific T cells. Nat Immunol 20, 1244–1255. 10.1038/s41590-019-0465-3.

36. Andreatta, M., and Carmona, S.J. (2021). UCell: Robust and scalable single-cell gene signature scoring. Comput Struct Biotechnol J 19, 3796–3798. 10.1016/j.csbj.2021.06.043.

37. Bugaut, H., El Morr, Y., Mestdagh, M., Darbois, A., Paiva, R.A., Salou, M., Perrin, L., Furstenheim, M., du Halgouet, A., Bilonda-Mutala, L., et al. (2024). A conserved transcriptional program for MAIT cells across mammalian evolution. J Exp Med 221. 10.1084/jem.20231487.

38. Hollbacher, B., Duhen, T., Motley, S., Klicznik, M.M., Gratz, I.K., and Campbell, D.J. (2020). Transcriptomic Profiling of Human Effector and Regulatory T Cell Subsets Identifies Predictive Population Signatures. Immunohorizons 4, 585–596. 10.4049/immunohorizons.2000037.

39. Pawlak, M., DeTomaso, D., Schnell, A., Meyer Zu Horste, G., Lee, Y., Nyman, J., Dionne, D., Regan, B.M.L., Singh, V., Delorey, T., et al. (2022). Induction of a colitogenic phenotype in Th1-like cells depends on interleukin-23 receptor signaling. Immunity 55, 1663–1679 e1666. 10.1016/j.immuni.2022.08.007.

40. Cantini, L., Zakeri, P., Hernandez, C., Naldi, A., Thieffry, D., Remy, E., and Baudot, A. (2021). Benchmarking joint multi-omics dimensionality reduction approaches for the study of cancer. Nat Commun 12, 124. 10.1038/s41467-020-20430-7.

41. Huizing, G.J., Deutschmann, I.M., Peyre, G., and Cantini, L. (2023). Paired single-cell multi-omics data integration with Mowgli. Nat Commun 14, 7711. 10.1038/s41467-023-43019-2.

42. McGonagle, D., and McDermott, M.F. (2006). A proposed classification of the immunological diseases. PLoS Med 3, e297. 10.1371/journal.pmed.0030297.

43. Mouri, K., Guo, M.H., de Boer, C.G., Lissner, M.M., Harten, I.A., Newby, G.A., DeBerg, H.A., Platt, W.F., Gentili, M., Liu, D.R., et al. (2022). Prioritization of autoimmune disease-associated genetic variants that perturb regulatory element activity in T cells. Nat Genet 54, 603–612. 10.1038/s41588-022-01056-5.

44. Utzschneider, D.T., Gabriel, S.S., Chisanga, D., Gloury, R., Gubser, P.M., Vasanthakumar, A., Shi, W., and Kallies, A. (2020). Early precursor T cells establish and propagate T cell exhaustion in chronic infection. Nat Immunol 21, 1256–1266. 10.1038/s41590-020-0760-z.

45. Yao, C., Lou, G., Sun, H.W., Zhu, Z., Sun, Y., Chen, Z., Chauss, D., Moseman, E.A., Cheng, J., D’Antonio, M.A., et al. (2021). BACH2 enforces the transcriptional and epigenetic programs of stem-like CD8(+) T cells. Nat Immunol 22, 370–380. 10.1038/s41590-021-00868-7.

46. Park, S.H., Rhee, J., Kim, S.K., Kang, J.A., Kwak, J.S., Son, Y.O., Choi, W.S., Park, S.G., and Chun, J.S. (2018). BATF regulates collagen-induced arthritis by regulating T helper cell differentiation. Arthritis Res Ther 20, 161. 10.1186/s13075-018-1658-0.

47. Punkenburg, E., Vogler, T., Buttner, M., Amann, K., Waldner, M., Atreya, R., Abendroth, B., Mudter, J., Merkel, S., Gallmeier, E., et al. (2016). Batf-dependent Th17 cells critically regulate IL-23 driven colitis-associated colon cancer. Gut 65, 1139–1150. 10.1136/gutjnl-2014-308227.

48. Schraml, B.U., Hildner, K., Ise, W., Lee, W.L., Smith, W.A., Solomon, B., Sahota, G., Sim, J., Mukasa, R., Cemerski, S., et al. (2009). The AP-1 transcription factor Batf controls T(H)17 differentiation. Nature 460, 405–409. 10.1038/nature08114.

49. Hasan, Z., Koizumi, S.I., Sasaki, D., Yamada, H., Arakaki, N., Fujihara, Y., Okitsu, S., Shirahata, H., and Ishikawa, H. (2017). JunB is essential for IL-23-dependent pathogenicity of Th17 cells. Nat Commun 8, 15628. 10.1038/ncomms15628.

50. Bentsen, M., Goymann, P., Schultheis, H., Klee, K., Petrova, A., Wiegandt, R., Fust, A., Preussner, J., Kuenne, C., Braun, T., et al. (2020). ATAC-seq footprinting unravels kinetics of transcription factor binding during zygotic genome activation. Nat Commun 11, 4267. 10.1038/s41467-020-18035-1.

51. Teng, M.W., Bowman, E.P., McElwee, J.J., Smyth, M.J., Casanova, J.L., Cooper, A.M., and Cua, D.J. (2015). IL-12 and IL-23 cytokines: from discovery to targeted therapies for immune-mediated inflammatory diseases. Nature medicine 21, 719–729. 10.1038/nm.3895.

52. Aggarwal, S., Ghilardi, N., Xie, M.H., de Sauvage, F.J., and Gurney, A.L. (2003). Interleukin-23 promotes a distinct CD4 T cell activation state characterized by the production of interleukin-17. The Journal of biological chemistry 278, 1910–1914. 10.1074/jbc.M207577200.

53. Wang, H., Kjer-Nielsen, L., Shi, M., D’Souza, C., Pediongco, T.J., Cao, H., Kostenko, L., Lim, X.Y., Eckle, S.B.G., Meehan, B.S., et al. (2019). IL-23 costimulates antigen-specific MAIT cell activation and enables vaccination against bacterial infection. Sci Immunol 4. 10.1126/sciimmunol.aaw0402.

54. Rahimpour, A., Koay, H.F., Enders, A., Clanchy, R., Eckle, S.B., Meehan, B., Chen, Z., Whittle, B., Liu, L., Fairlie, D.P., et al. (2015). Identification of phenotypically and functionally heterogeneous mouse mucosal-associated invariant T cells using MR1 tetramers. J Exp Med 212, 1095–1108. 10.1084/jem.20142110.

55. Koay, H.F., Gherardin, N.A., Enders, A., Loh, L., Mackay, L.K., Almeida, C.F., Russ, B.E., Nold-Petry, C.A., Nold, M.F., Bedoui, S., et al. (2016). A three-stage intrathymic development pathway for the mucosal-associated invariant T cell lineage. Nat Immunol 17, 1300–1311. 10.1038/ni.3565.

56. Chandra, S., Ascui, G., Riffelmacher, T., Chawla, A., Ramirez-Suastegui, C., Castelan, V.C., Seumois, G., Simon, H., Murray, M.P., Seo, G.Y., et al. (2023). Transcriptomes and metabolism define mouse and human MAIT cell populations. Sci Immunol 8, eabn8531. 10.1126/sciimmunol.abn8531.

57. Dott, T., Culina, S., Chemali, R., Mansour, C.A., Dubois, F., Jagla, B., Doisne, J.M., Rogge, L., Huetz, F., Jonsson, F., et al. (2024). Standardized high-dimensional spectral cytometry protocol and panels for whole blood immune phenotyping in clinical and translational studies. Cytometry A 105, 124–138. 10.1002/cyto.a.24801.

58. Ritchie, M.E., Phipson, B., Wu, D., Hu, Y., Law, C.W., Shi, W., and Smyth, G.K. (2015). limma powers differential expression analyses for RNA-sequencing and microarray studies. Nucleic Acids Res 43, e47. 10.1093/nar/gkv007.

59. Liao, Y., Wang, J., Jaehnig, E.J., Shi, Z., and Zhang, B. (2019). WebGestalt 2019: gene set analysis toolkit with revamped UIs and APIs. Nucleic Acids Res 47, W199–W205. 10.1093/nar/gkz401.

60. Amemiya, H.M., Kundaje, A., and Boyle, A.P. (2019). The ENCODE Blacklist: Identification of Problematic Regions of the Genome. Sci Rep 9, 9354. 10.1038/s41598-019-45839-z.

61. Corces, M.R., Granja, J.M., Shams, S., Louie, B.H., Seoane, J.A., Zhou, W., Silva, T.C., Groeneveld, C., Wong, C.K., Cho, S.W., et al. (2018). The chromatin accessibility landscape of primary human cancers. Science 362. 10.1126/science.aav1898.

62. Labun, K., Montague, T.G., Krause, M., Torres Cleuren, Y.N., Tjeldnes, H., and Valen, E. (2019). CHOPCHOP v3: expanding the CRISPR web toolbox beyond genome editing. Nucleic Acids Res 47, W171–W174. 10.1093/nar/gkz365.

63. Menegatti, S., Guillemot, V., Latis, E., Yahia-Cherbal, H., Mittermuller, D., Rouilly, V., Mascia, E., Rosine, N., Koturan, S., Millot, G.A., et al. (2021). Immune response profiling of patients with spondyloarthritis reveals signalling networks mediating TNF-blocker function in vivo. Ann Rheum Dis 80, 475–486. 10.1136/annrheumdis-2020-218304.

