## Supplemental Figures for "IL-23 tunes inflammatory functions of human mucosal-associated invariant T (MAIT) cells"

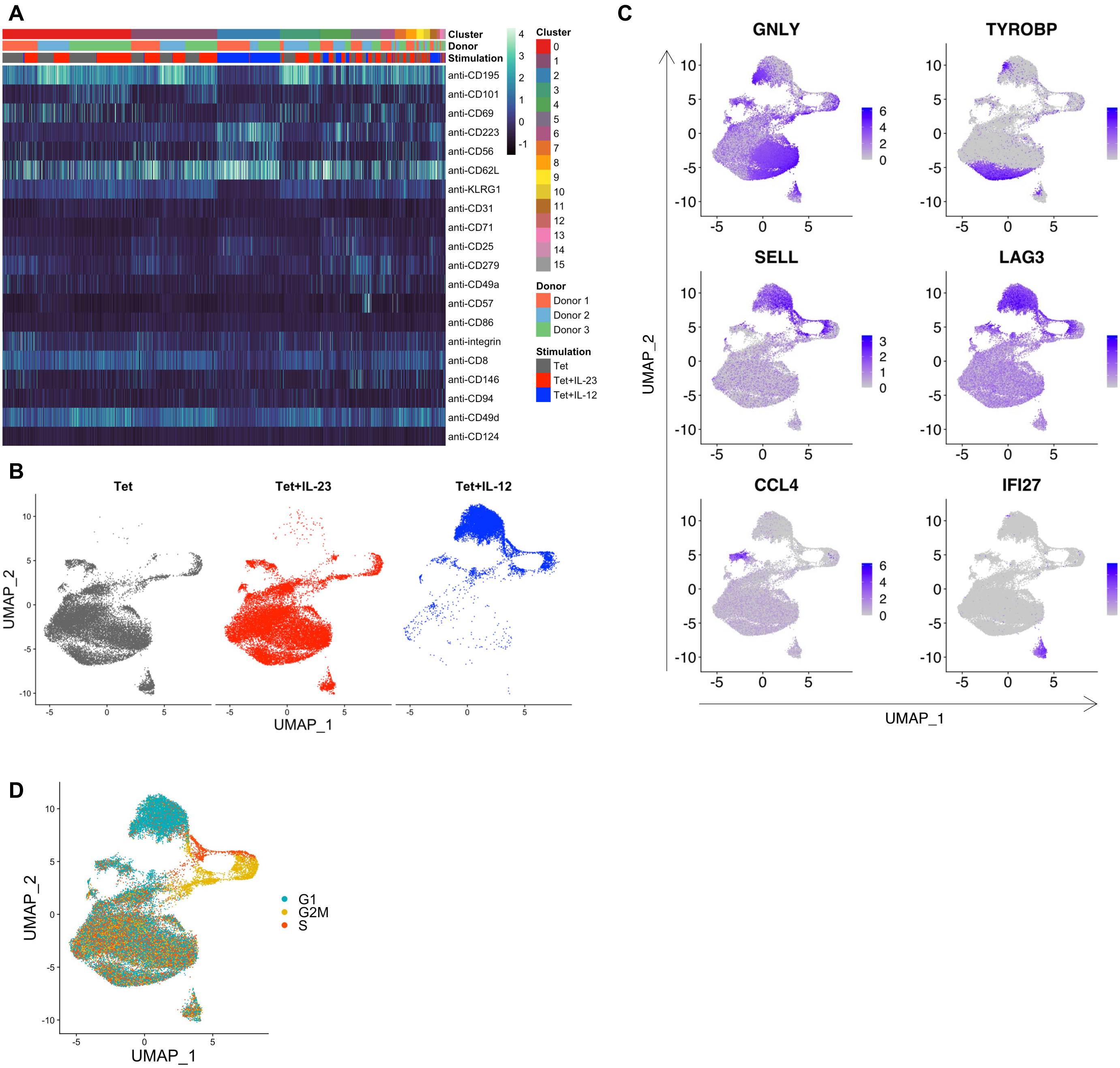


**Fig. S1. Single-cell phenotypic and transcriptional analysis of stimulated MAIT cells.**

(**A**) Heatmap showing row-scaled normalized expression of the top 3 antibody markers for each MAIT cell cluster. (**B**): Projection on the UMAP of cells according to their stimulation group. (**C**) Projection on the UMAP of the expression of selected cluster marker genes. (**D**) Projection on the UMAP of the cell cycle phase.

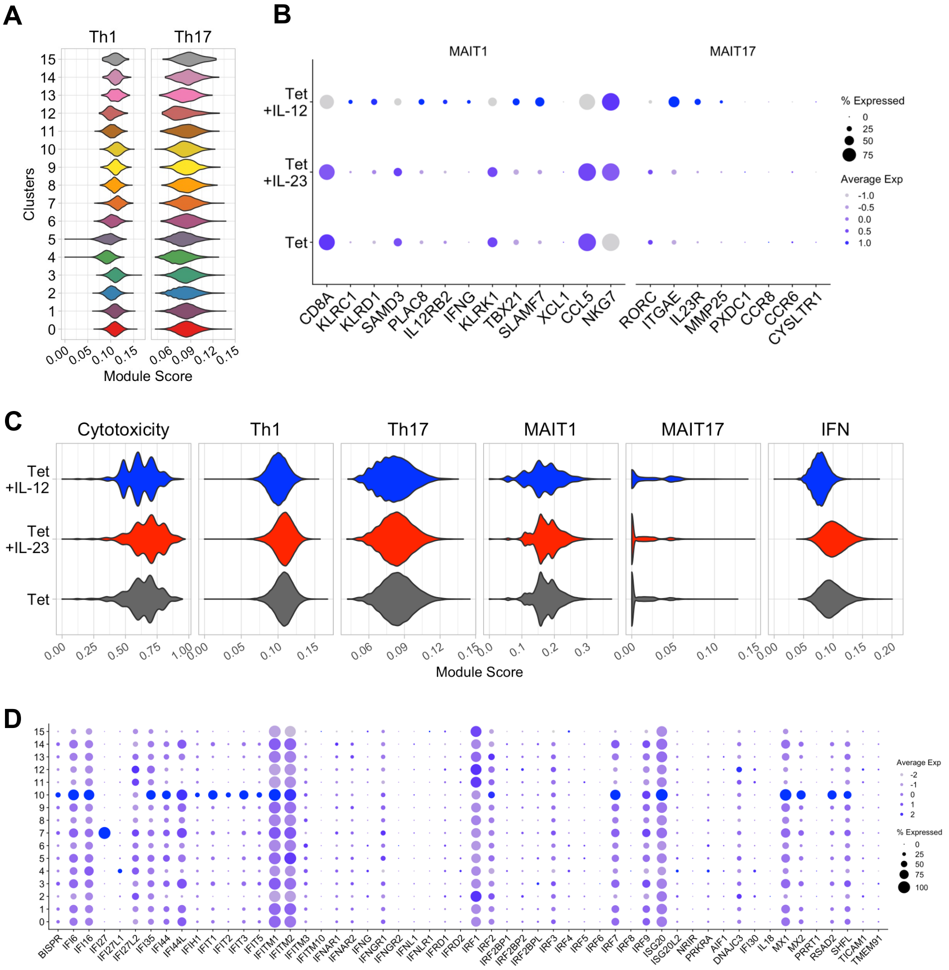


**Fig. S2. Functional characterization of stimulated MAIT cells.**

(**A**) Violin plots of Th1 and Th17 signature scores by cluster. (**B**) Dot plot of MAIT1 and MAIT17 signature genes by stimulation group. (**C**) Violin plots of the indicated signatures by stimulation group. (**D**) Dot plot of IFN signature genes by cluster.

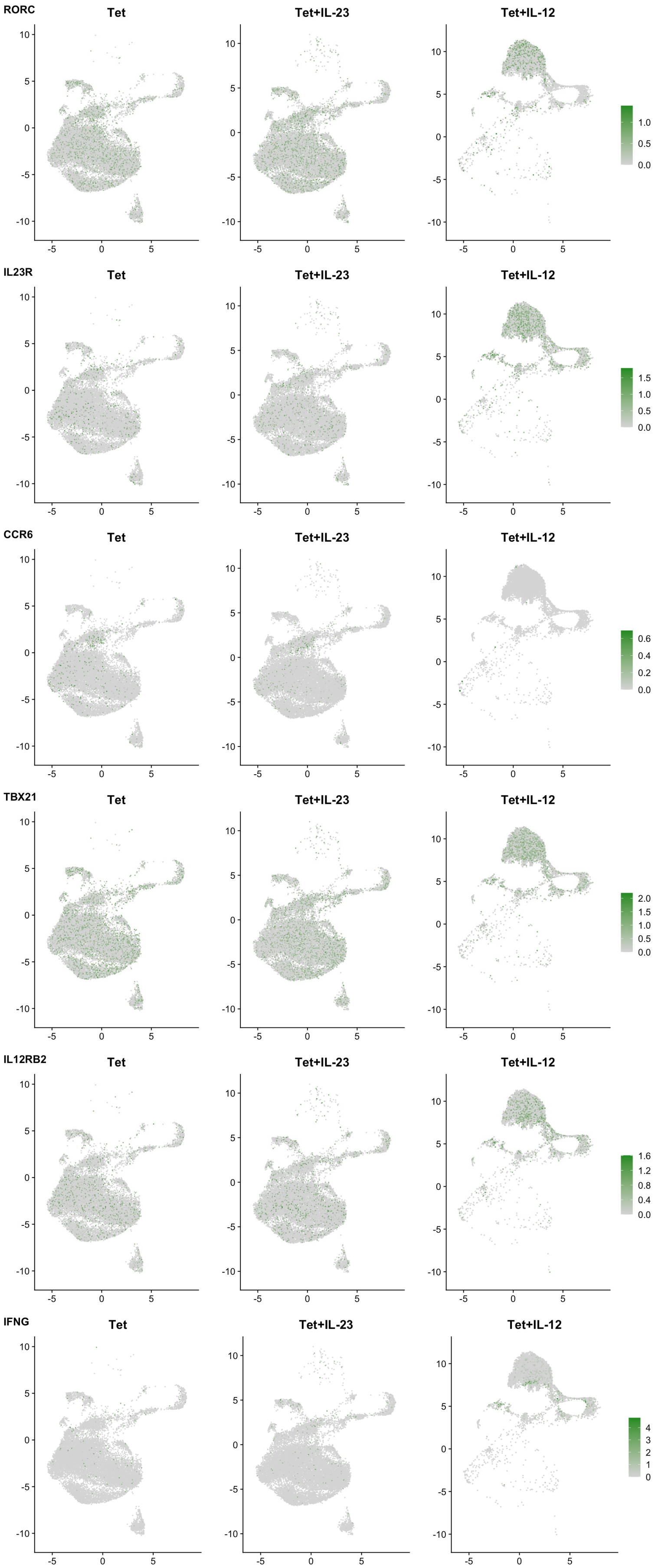


**Fig. S3. Expression of type 17 and type 1 marker genes at the single-cell level.**

The expression of type 17 (*RORC, IL23R, CCR6*) and type 1 (*TBX21, IL12RB2, IFNG*) responses genes was projected on the UMAPs representing the subset of MAIT cells stimulated in the indicated condition.

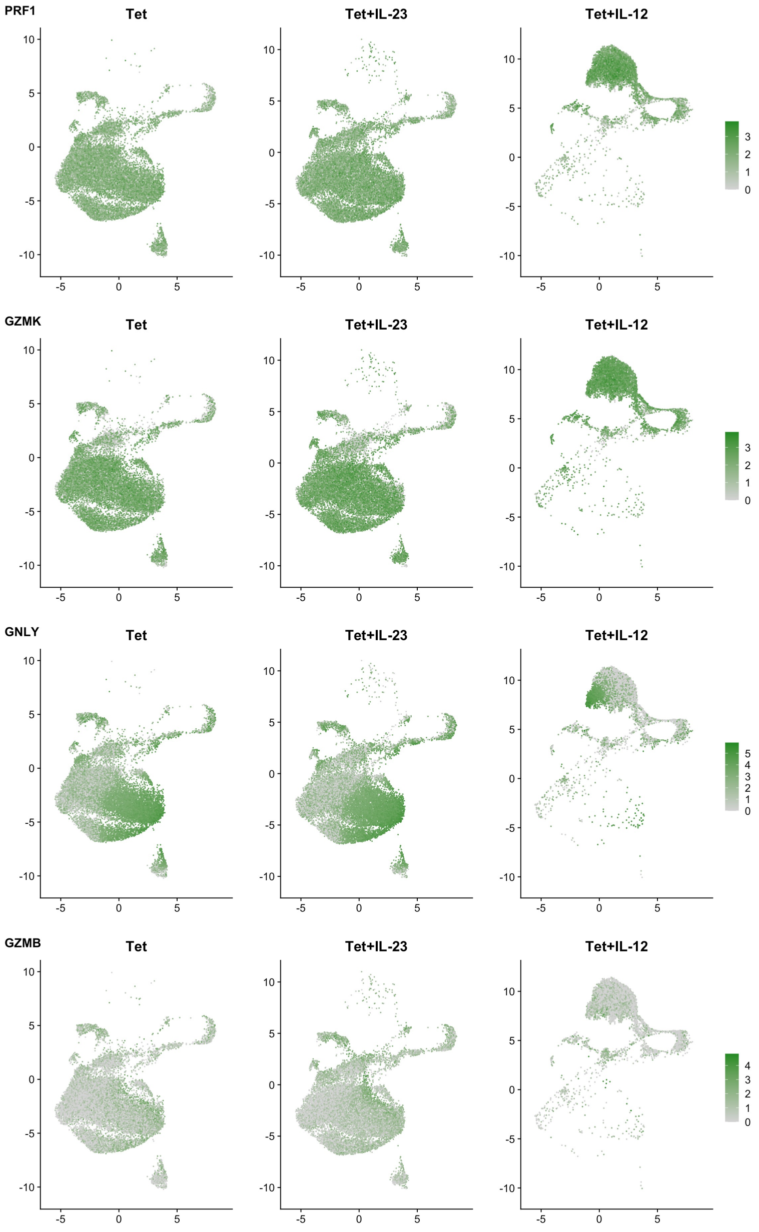


**Fig. S4. Expression of genes encoding cytotoxic molecules at the single-cell level.**

The expression of genes encoding cytotoxic molecules was projected on the UMAPs representing the subset of MAIT cells stimulated in the indicated condition.

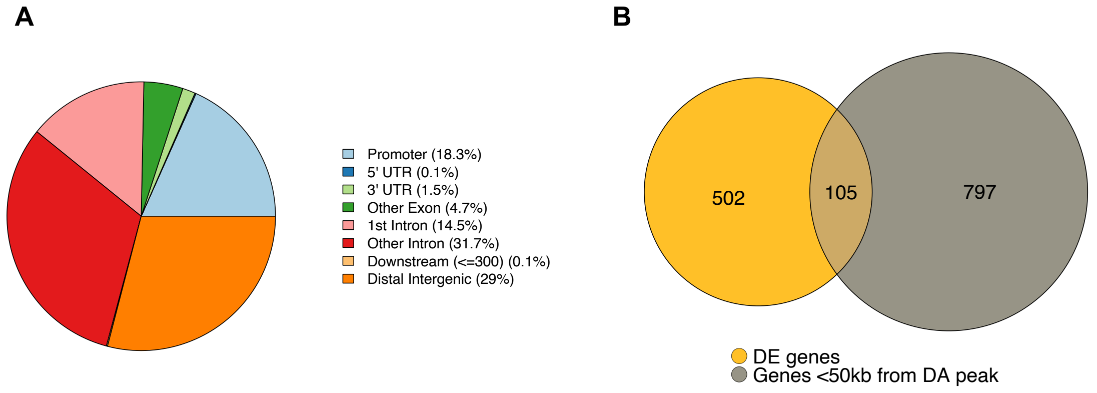


**Fig. S5. Characteristics of differentially accessible regions and intersection with differentially expressed genes.**
(**A**) Distribution of differentially accessible (DA) regions (Tet+IL-23 vs. Tet) across the main gene-related annotation categories. (**B**) Venn diagram displaying the overlap between differentially expressed genes and genes in the vicinity (<50kb from TSS) of a DA region.
